# Honey bee (*Apis mellifera*) gut microbiome associations with viruses and pesticides across Canadian agroecosystems

**DOI:** 10.64898/2026.06.23.731697

**Authors:** Aleksandra Kozlova-Ryabova, Lan Tran, Lance Lansing, Morgan Cunningham, Jonathan Ho, Thomas B Deckers, Amanda S. Gregoris, Jackie Zorz, Sarah K French, Aidan Jamieson, Mateus Pepinelli, Ida M Conflitti, Pierre Giovenazzo, Shelley E. Hoover, Robert W. Currie, Stephen F. Pernal, Amro Zayed, Rodrigo Ortega Polo, Hosna Jabbari, M. Marta Guarna, Leonard J. Foster, Huan Zhong

## Abstract

The honey bee (*Apis mellifera*) gut microbiome plays a central role in host health, yet its variation across agricultural landscapes remains poorly resolved. This study investigates how major environmental stressors, particularly pesticide exposure and RNA virus loadings, shape the honey bee gut microbiome in a large-scale field study conducted across Canada, spanning diverse agroecosystems from British Columbia to Quebec. We identify consistent associations between specific bacterial taxa and major RNA viruses, including enrichment of *Serratia marcescens* with SBV and depletion of *Bombella intestini* with BQCV. Pesticide exposure is likewise linked to reproducible shifts in key microbial taxa. Together, these findings reveal that interacting stressors jointly shape the bee gut microbiome and enable prediction of microbiome responses in agroecosystems.

**Highlights:**

Distinct associations identified between gut bacteria and major bee RNA viruses (BQCV, SBV, LSV, IAPV)
Pesticide exposure is linked to reproducible shifts in key microbial taxa
Combined virus–pesticide effects form coordinated clusters that predict microbiome variation and specific bacterial responses
Integrated modeling demonstrates that environmental stressors can jointly explain microbiome structure beyond crop effects

**Graphical abstract:** Schematic overview of potential links between pesticide exposure and RNA virus infection and their effects on the bee gut bacterial community. Solid arrows indicate associations supported by the present study, whereas dashed arrows indicate hypothesized or unresolved interactions. Associations between the presence of specific bee RNA viruses (left) or pesticide residues (right) and changes in the relative abundance of particular gut taxa (pink ↑, increased; blue ↓, decreased). The pesticide subtype is indicated by the icon in the cell (leaf - herbicide, hyphae - fungicide and insect - insecticide).

Several bacterial taxa showed reproducible associations with specific viral or pesticide variables, including *Bombella intestini, Serratia marcescens, Melissococcus plutonius, Paenibacillus alvei, Apibacter* sp. wkB309, and *Gilliamella* sp. A7.

Abbreviations: BQCV Black queen cell virus; LSV, Lake Sinai virus; SBV, Sacbrood virus; IAPV, Israeli acute paralysis virus. (p/n/b) indicate the sample matrix in which the pesticide was detected, namely pollen, nectar, and bee tissue, respectively.

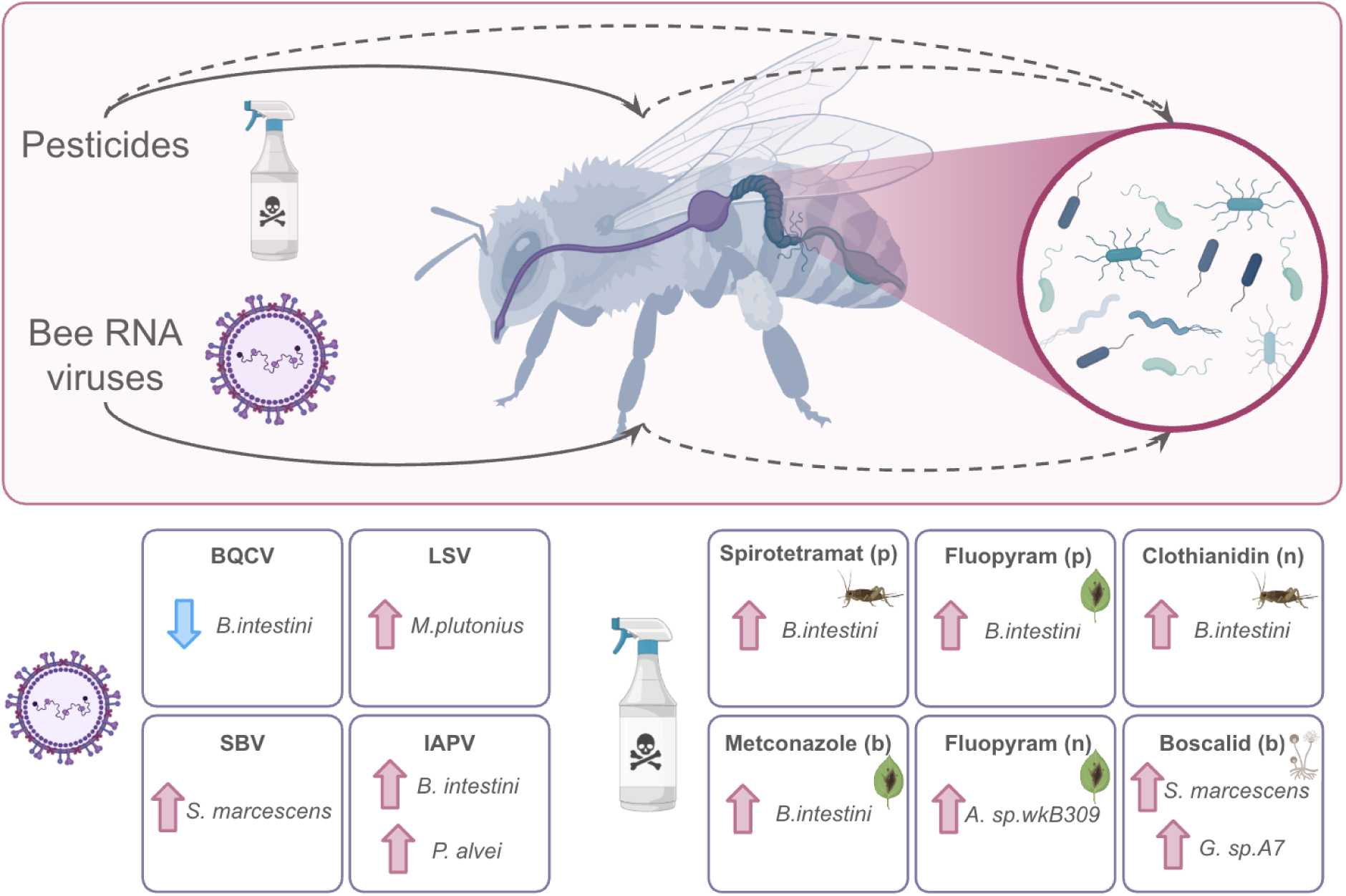

## Introduction

The Western honey bee (*Apis mellifera*) is a vital pollinator in global ecosystems and agriculture^1–4^. Unfortunately, substantial evidence points to widespread problems connected to bee populations^5,6^. There is evidence that annual winter mortality is increasing and is not consistent with yearly likely colony recovery, and the global population of domesticated honeybees is growing more slowly than the demand for pollination^7–12^.

Honey bee health is undermined by several interrelated stressors, including intensive pesticide use, and pathogen and parasite infections^13–15^. These effects can be collinear and difficult to separate with single-factor analysis, necessitating multi-factor statistical models. Given the multitude of environmental stressors bees face, it has become increasingly clear that their resilience depends not only on external conditions but also on internal biological systems. One of the most critical yet often overlooked components of bee health is the composition of their gut microbiome^16,17^. This complex community of symbiotic bacteria plays a pivotal role in digestion, immunity, detoxification, and development^18^. Disruptions to the microbiome, whether due to pesticides, poor nutrition, or pathogen exposure, can weaken colony health and amplify the effects of other stress factors^19,20^. Understanding the structure and function of the bee gut microbiota is therefore essential for developing strategies to support pollinator health in changing environments.

The gut microbiome of *A. mellifera* is a relatively conserved community dominated by a limited set of core bacterial genera that are consistently detected across individuals, colonies, and bee populations, together with a more variable set of non-core taxa^21,22^. Core members of this community typically include *Gilliamella*, *Snodgrassella*, *Bifidobacterium*, *Lactobacillus*-related lineages, *Frischella*, *Bartonella*, and *Commensalibacter*, many of which contribute to carbohydrate metabolism, immune modulation, and resistance to pathogens^21–23^.

However, variations in this core composition are driven by diet, habitat, or disease, and can significantly affect bee resilience^21,24–26^. Despite substantial progress in describing the structure and function of the honey bee gut microbiome, it remains unclear how microbiome variation is structured across real-world agricultural environments in which crop type, pathogen burden, and pesticide exposure varies. In particular, only a few studies by us and others have addressed how microbiome composition responds to variation in pathogen and pesticide exposure^24,27–32^.

The goal of this study is to provide a comprehensive analysis of the gut microbiota of Canadian *A. mellifera* colonies and their association with pesticides and viruses. We aim to characterize the overall composition of the core microbiome and assess the variability of microbial communities across samples from different crop ecosystems (1), and investigate the abundance of known immunostimulatory symbionts in relation to environmental parameters, such as pesticides (2) and pathogens (3). By integrating microbiological profiling with ecological metadata, this work contributes to a better understanding of the local dynamics of bee-microbe interactions and their implications for colony health.

## Results

### Microbiome community composition

To conduct the experiment in 2020 and 2021, bee samples were collected from eight crop ecosystems in the ’mid-bloom’ for the crop: Commodity canola (CAC), Seed canola (CAS), Corn (COR), Cranberry (CRA), Highbush blueberry (HBB), Lowbush blueberry (LBB), Soybean (SOY), and Apple (APP) (Figure 1a)^33,34^. For each crop ecosystem, five to ten apiaries were selected near flowering plants associated with that crop ecosystem. Twelve bees were collected from each apiary, their ’guts’, i.e. whole digestive system except for the foregut/crop were isolated, and pooled into a single biological replicate (Figure 1b). DNA was extracted from these samples and shotgun sequencing was performed.

**Figure 1.**
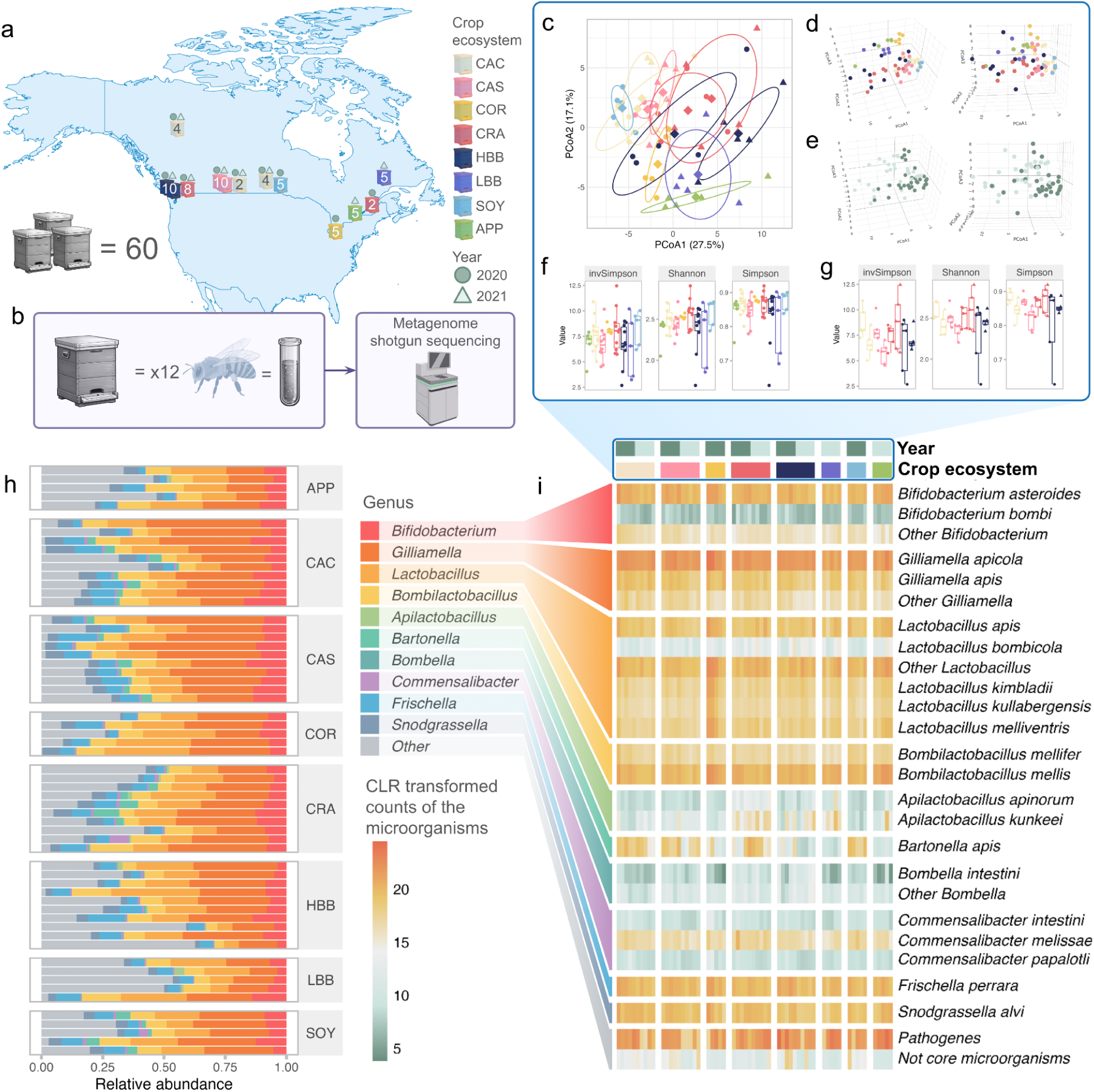
Microbiome dynamics across samples. **(a)** The map shows sampling points in the shape of hives. The hives are colored according to the crop ecosystems on which the bees were placed (CAC = Canola oil, CAS = Canola seed, COR = Corn, CRA = Cranberry, HBB = Highbush blueberry, LBB = Lowbush blueberry, SOY = Soybean, APP = Apple). The numbers indicate how many apiaries (samples) were sampled in each region. **(b)** Each group of 12 bee guts from an apiary (taken from four colonies) was combined into a single sample and sequenced on the NovaSeq Illumina platform. **(c)** A two-dimensional plot of the principal coordinate (PCoA) method using the Aitchison distance. The axes show % variance explained. The dots are colored according to the crop ecosystem. The shape of the dots corresponds to the year in which the samples were collected. The graph contains ellipses to represent groups of samples belonging to the same crop ecosystem. For samples belonging to the same crop ecosystem but collected in different years, two ellipses of the corresponding color are shown. **(d,e)** A three-dimensional plot of the principal coordinate (PCoA) method using the Aitchison distance. The dots are colored according to the (d) crop ecosystem and (e) year. The plot is shown from two different angles. **(f,g):** Alpha-diversity indices shown as boxplots with pairwise comparisons, grouped by crop ecosystem (f) and sampling year (g). Boxplots show medians, interquartile ranges, and whiskers extending to 1.5 × IQR. *P* values were adjusted using the Benjamini–Hochberg method. **(h)** Relative abundance of whole microbial community composition in the bee gut samples as the number of genome equivalents calculated with Kraken. Splitting on the groups was performed based on the crop environment. Strips are equal to the crop environment abbreviations. **(i)** Heatmap of the CLR-transformed microbial composition. Rows are species, grouped by genera and color-coded, columns are samples. Column annotation colors indicate sampling crop ecosystems and year and row annotation colors indicate microorganisms in connection to the group of microorganisms.

Microbiome profiling using the BeeRoLaMa v1 database successfully identified all major microbial groups known to inhabit the honey bee gut, including protists, fungi, and bacteria. In total, 55 microbial taxa were identified across all samples. In each sample, an average of 72.3% (maximum 79.2%) of the reads mapped to the host genome, while the remaining 27.7% reads were assigned to bacterial taxa. Scaling the non-host reads to 100%, an average of 12.3% of reads remained unclassified, with a range of 8.9% to 32.1%. All tables containing microbial abundance data for each sample are available on GitHub: (https://github.com/andwhoami/BeeCsiMetagenome).

Across samples, the gut microbiome was dominated by canonical honey bee-associated taxa, including *Gilliamella*, *Snodgrassella*, *Frischella*, *Bartonella*, *Lactobacillus*-related lineages, and *Bifidobacterium* (Figure 1h). Overall, the microbiome composition matched the expected bacterial community typically found in the honey bee gut. The relative abundance of these core genera was largely consistent with previously reported bee microbiomes, where *Gilliamella* and *Snodgrassella* dominated, particularly in bees sampled between May and July^21^. Lower levels of *Lactobacillus* and *Bifidobacterium* further supported these expected microbial distributions.

To analyze microbial abundance patterns appropriately, we transformed the compositional microbiome data using a centered log-ratio (CLR)^35^ approach, which mitigates compositional constraints and enables Euclidean-based statistical analysis (Figure 1i). We derived a heatmap for the transformed CLR data of bacterial organisms at the species level. Essentially, we observed all the standard bacterial species in the bee’s gut. Representatives of *Lactobacillus* included the expected major and minor species, such as *L. kunkeei* which is characteristic of the crop and the hindgut but it is also found in the midgut34. In general, the distribution across the samples was quite homogeneous and had no clearly dominant genera.

To determine whether the crop environment influences the honey bee gut microbiome composition, we assessed differences in microbial communities sampling across crop ecosystems. We tested the spread of variances within the crop ecosystems using the betadisper function from the vegan package based on Aitchison distance, which showed significant differences (*p* < 0.001, F = 5.1289). Principal Coordinate Analysis (PCoA) (Figures 1c,d) supported this result, showing distinct clustering of microbial communities according to the crop type.

We performed the same process to compare samples collected in 2020 and 2021 (Figure 1e). Visual clustering of samples taken in different years can be explained by the different composition of crops within each year; nevertheless there were statistically significant differences only for microbiome composition (betadisper p = 0.21 and adonis < 0.01). To evaluate year effects without confounding by crop-specific sampling structure, we restricted the comparison to crop ecosystems represented in both years. We found that only in the CAS crop ecosystem was there a significantly different dispersion in the microbiome between samples collected in different years (adonis *p* = 0.007) (Supplementary note 1).

To further characterize how crop environments might influence the honey bee gut microbiome, we also assessed within-sample microbial diversity (alpha-diversity). We calculated the Simpson, Shannon, and Chao1 indices to estimate richness and evenness. The hypothesis that bees from different pollination environments would exhibit differences in alpha-diversity was not supported by the Wilcoxon test (p > 0.05 for all indices) (Figure 1f), indicating that the crop type did not affect the overall microbial diversity within individual bees.

To identify microbial drivers of the observed beta-diversity differences, we performed differential abundance analysis with ANCOM-BC2^36^. This analysis revealed taxa with significant shifts in abundance across crop ecosystems (adjusted p < 0.05, Supplementary note 2). In total, 33 of 55 bacterial species demonstrated differential representation in at least one crop environment, indicating that specific microbial lineages contribute to ecosystem-associated variation in the honey bee gut microbiome.

Given that agricultural environments vary not only in crop species but also in broader ecological and anthropogenic factors, we further partitioned the crop ecosystem into contributing components, including flower representation, virus burden, and pesticide exposure (Supplementary note 3). These metadata variables were examined in relation to the crop type and subsequently tested for associations with microbiome composition to identify which environmental factors most strongly influence gut microbial structure (Figure 2a, 3a).

**Figure 2.**
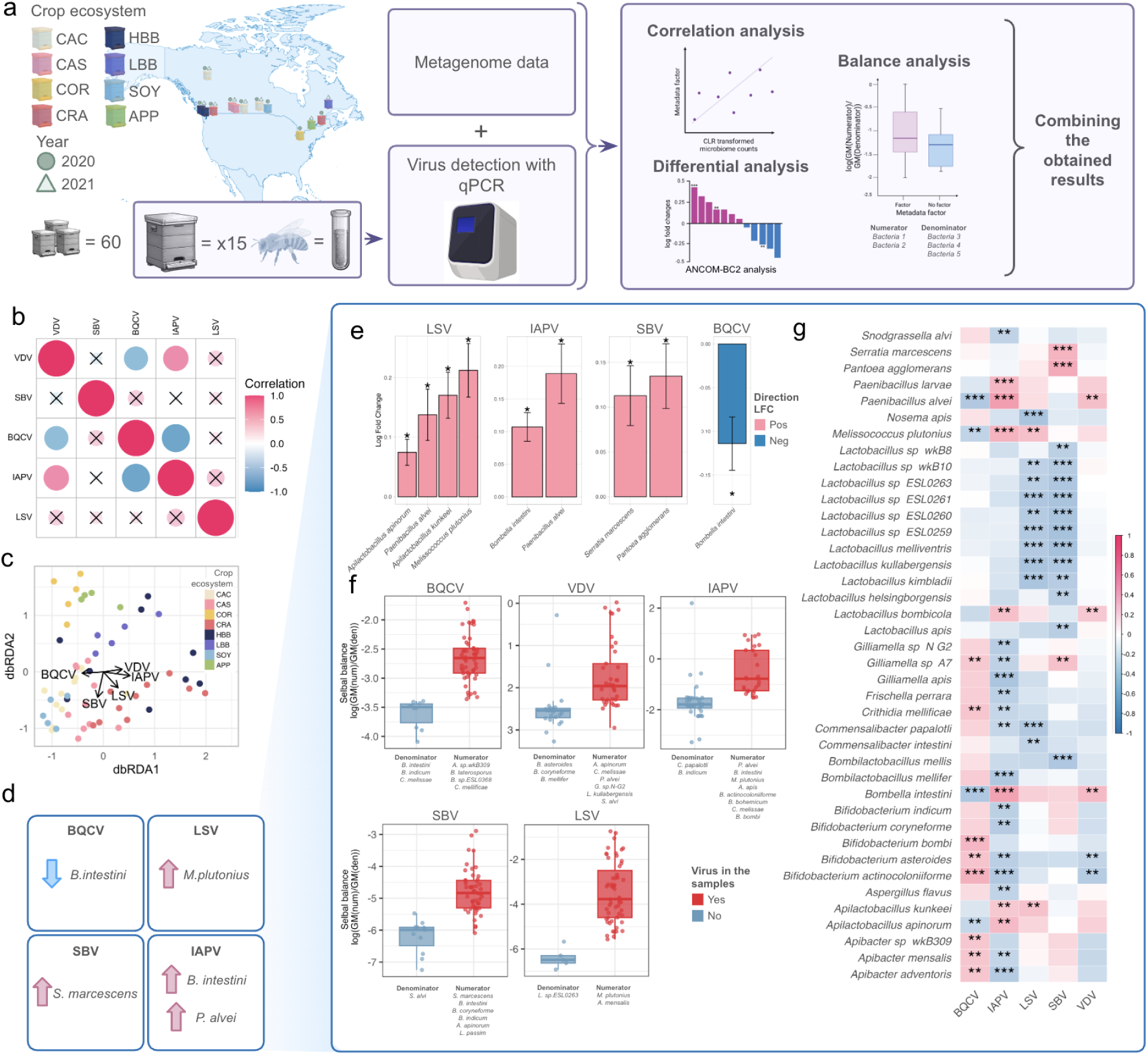
Microbiome connection to the bee viruses. **(a)** Sampling bees and processing samples for bee RNA virus detection with qPCR as described in McAfee *et al.* (2024)^39^ and analysis of the relationship between microbial organisms and viral loads. The analysis was performed using three methods and pooling of results. **(b)** Correlation plot for virus variables with each other based on the Spearman index. “X” indicates insignificant correlation. **(c)** Fdb-RDA ordination plots display microbial community composition constrained by virus variables. Vectors are targeted at crops in which viruses are present in high numbers. **(d)** Summary of microbiome associations with viruses **(e)** Log-fold change (LFC) of species abundance in the absence and presence of viruses across all samples and years. Statistical significance (ANCOM-BC2; p <0.05) is indicated by “*”. Positive LFC indicates an increase in abundance in the presence of a virus. **(f)** Each boxplot shows the SelBal balance comparing samples in which the virus is absent (“No”) versus present (“Yes”). If the “Yes” group displays higher balance values than the “No” group, this indicates that the numerator taxa are relatively more abundant (or the denominator taxa less abundant) in virus-positive samples. Individual points represent single samples. Statistical significance was assessed using a two-group comparison test (Wilcoxon rank-sum test), and the corresponding *p* value is shown. **(g)** Correlation heatmap between the virus representation and CLR-transformed microbe representation. Cells show the Spearman rank correlations (ρ) calculated across samples between each microbial trait (rows; CLR-transformed abundance) and each viral variable (columns; log-transformed, e.g., log10(x*107+1)). The color scale encodes ρ (blue = negative, red = positive; intensity = magnitude). Asterisks denote FDR-corrected significance (Benjamini–Hochberg for all pairs tested): q < 0.05 (*), q < 0.01 (**), q < 0.001 (***). Only traits that passed the prevalence/abundance filters are shown.

### Viruses in connection to crop ecosystem and microbiome composition

To investigate the associations between viral burden and gut microbiome composition, we focused on five viruses that passed prevalence filtering, including Black Queen Cell Virus (BQCV), Sacbrood Virus (SBV), Deformed Wing Virus - B (DWV-B previously known as Varroa Destructor Virus-1, VDV-1), Lake Sinai Virus (LSV), and Israeli Acute Paralysis Virus (IAPV), collected in 2020 and 2021 in the ’mid-bloom’ for the crop at the same collection points as the metagenomic samples. To characterize the relationships between each of them, correlation analysis, Distance-based ReDundancy Analysis (db-RDA), and collinearity testing were conducted (Figure 2b,c). This revealed a statistically significant relationship between some of the viruses (Figure 2b), allowing us to determine the combinations included in the analysis (Supplementary note 4).

After thoroughly characterizing virus variables, we conducted several complementary analyses to investigate how viral infection profiles relate to gut microbiome composition. Based on the virus screening results, we selected variables to include in microbiome–virus association models (Supplementary note 5). Because we found a strong link between environmental parameters and the сrop ecosystem, and considering that ANCOM-BC2 and SelBal rely on regression models that are highly sensitive to correlated predictors, we conducted subsequent analyses in several different ways: without adjusting for the crop ecosystem, with adjusting for the crop ecosystem, and individually for several separated crop ecosystems to confirm the observed relationship without adjusting for this parameter (Supplementary note 6, Supplementary figures 1-5)^37,38^.

In the ANCOM-BC2 analysis performed without crop adjustment, we identified bacterial taxa significantly associated with four of the five viruses examined (Figure 2e). For LSV, IAPV, and SBV, multiple bacterial species showed increased abundance in infected bees, indicating positive associations with viral presence. Conversely, *Bombella intestini* was more abundant in samples without BQCV infection, consistent with a negative association between this bacterium and the virus.

For correlation analysis based on CLR transformation with each virus, statistically significant connected microorganisms for both Spearman and Pearson correlation were found (Figure 2g). For *B. intestini* and *P. alvei*, significant positive correlation was found with IAPV and VDV. Additionally, IAPV and VDV were positively significantly correlated with each other. BQCV virus was found to have a negative correlation with the same bacteria.

In the SelBal analysis, we constructed models without the crop-ecosystem correction and evaluated each virus independently (Figure 2f)^40^. *Commensalibacter melissae* and *Paenibacillus alvei* consistently appeared in the numerator of balances associated with IAPV and VDV, indicating increased abundance in bees infected with these viruses. In contrast, *C. melissae* and *B. intestini* formed a part of the denominator in the BQCV-associated balance. These patterns align with the correlation results, again demonstrating negative associations between BQCV and these bacterial taxa and positive associations with IAPV and VDV.

To find microorganisms that were more likely to be associated with the presence of the virus, we made an intersection of all performed analysis. As a result, we identified several bacterial species, which had significant, consistent tendencies for each analysis method (Table 1, Figure 2d). For LSV, *M. plutonius* was observed to be significantly more abundant in samples with higher virus load. In all three types of analysis, *B. intestini* and *P. alvei* were also found to be significantly associated with IAPV. For SBV, significantly higher abundance of *S.marcescens* was discovered, while for BQCV, there was a significantly lower level of *B.intestini*. For VDV, no associations with taxa were found.

**Table 1.** Microorganisms common for different types of analysis. The rows show combinations of the analyses. The columns correspond to viruses, and each cell shows bacteria whose abundance was positively or negatively correlated with that of the virus. For SelBal, a microorganism was classified as positively associated if it was present in the “numerator” group and negatively associated, if it was in the “denominator” group.

| Virus | LSV |  | IAPV |  | SBV |  | BQCV |  | VDV |  |
| --- | --- | --- | --- | --- | --- | --- | --- | --- | --- | --- |
| Effect upon exposure | Positive | Negative | Positive | Negative | Positive | Negative | Positive | Negative | Positive | Negative |
| ANCOM-BC<br>2 x SelBal | *M. plutonius* | - | *B. intestini*<br>*P. alvei* | - | *S. marcescens* | - | - | *B. intestini* | - | - |
| ANCOM-BC<br>2 x<br>Correlations<br>(clr) | *A. kunkeei*<br>*M. plutonius* | - | *B. intestini*<br>*P. alvei* | - | *S. marcescens*<br>*P. agglomerans* | - | - | *B. intestini* | - | - |
| Correlations<br>(clr) x SelBal | *M. plutonius* | *L. sp.ESL0* | *B. intestini*<br>*P. alvei* | *C. papalotli* | *S. marcescens* | - | *A. sp.wkB* | *B. intestini* | *P. alvei* | *B. asteroides* |
|  |  | 263 | *M. plutonius* | *B. indicum* |  |  | 309 |  |  |  |
| **General species** | *M. plutonius* | - | *B. intestini*<br>*P. alvei* | - | *S. marcescens* | - | - | *B. intestini* | - | - |
| General counts | 1 | 0 | 2 | 0 | 1 | 0 | 0 | 1 | 0 | 0 |

In general, based on the results, it is clear that 4 viruses are associated with 4 bacterial organisms, where 2 organisms are associated with 1 virus, and 1 organism is associated with 2 viruses. In all analysis options: 1) without adjusting for the crop ecosystem, 2) with adjusting for the crop ecosystem, and 3) individually for several separated crop ecosystems, we observed similar trends in the association of the microbiome with pesticide and virus parameters.

For a deeper explanation of the microbiome differences and in search for the connection between microbiome and the levels of pesticides, we applied the same analysis to the pesticide variables.

### Pesticides in connection to crop ecosystem and microbiome composition

To examine the association between pesticides and the microbiome, we used data on pesticides in nectar (n), pollen (p), and bees (b), collected in 2020 and 2021 in the ’mid-bloom’ for the crop at the same collection points as the metagenomic and viral samples. A total of 232 pesticides were initially assessed in these samples, but most were detected in only 2-3 samples out of 60. Since this is insufficient to identify a relationship between microbes and pesticides, we only included pesticides present in 30% of the samples^41,42^. As a result, nine pesticides were used: five were detected in pollen, two in nectar, and two in bee tissue.

As with the pathogen analyses, we first conducted a comprehensive characterization of variables before selecting those included in the microbiome–pesticide association models (Supplementary note 7) and used previously published results^43^. Collinearity testing and permutation analyses revealed substantial overlap between pesticide variables and crop ecosystem factors, underscoring the need for caution when incorporating both into regression-based frameworks. We therefore ran ANCOM-BC2 using multiple combinations of variables (Supplementary notes 8,9, Supplementary figures 8-16) and retained only microbial taxa that were consistently identified across models for primary reporting (Figure 3b). This conservative strategy ensured robust identification of microbiome features associated with pesticide exposure while accounting for ecological covariation.

**Figure 3.**
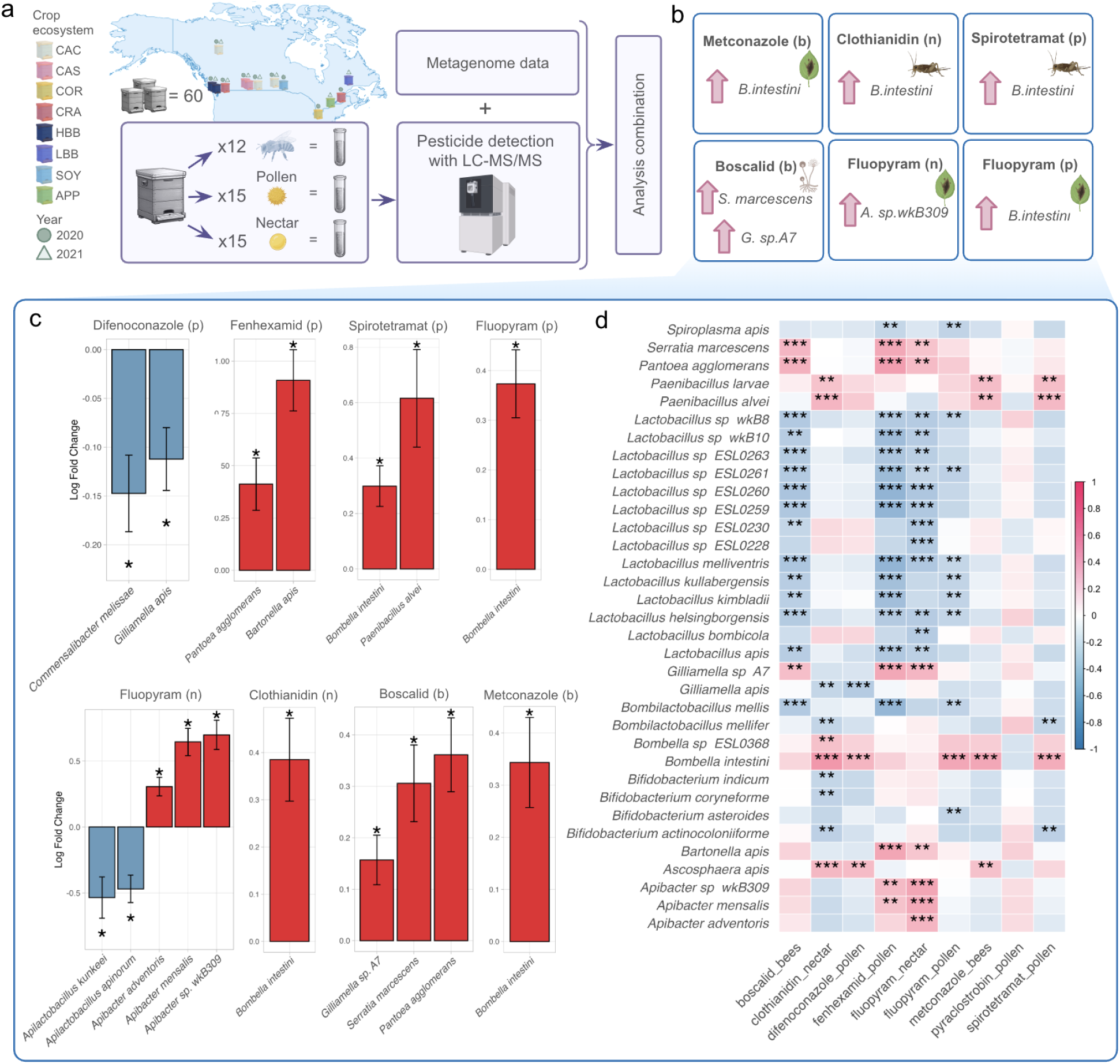
Microbiome connection to the pesticides. **(a)** The map shows sampling points. The points correspond to the locations where samples were collected for metagenomic and viral studies. From each point, 12 bee samples, 15 pollen samples, and 15 nectar samples were collected. The 12 bees were combined into one replicate. The same was done for the pollen and nectar samples. Pesticide analysis was then performed using LC-MC/MC. The resulting data were combined with microbial data and analyzed using three different methods. The results obtained for the three different methods were combined. **(b)** Microbiome association with pesticides. **(c)** Log fold change (LFC) of species abundance between pesticide present or absent from all samples and years. The plot used only statistically significant different bacteria. Statistical significance (ANCOM-BC2; p <0.05) is indicated by *. Positive LFC indicates an increase in abundance between factors, presence compared to absence for each pesticide, whereas negative LFC indicates a decrease. **(d)** Cells show Spearman rank correlations (ρ) calculated across samples between each microbial trait (rows; CLR-transformed abundance) and each pesticide variable (columns; log-transformed, e.g., log10(x*107+1)). The color scale encodes ρ (blue = negative, red = positive; intensity = magnitude). Asterisks denote FDR-corrected significance (Benjamini–Hochberg for all pairs tested): q < 0.05 (*), q < 0.01 (**), q < 0.001 (***). Only traits that passed the prevalence/abundance filters are shown.

In the ANCOM-BC2 analysis performed without crop adjustment, we identified bacterial taxa significantly associated with all pesticides (Figure 3c). *Bombella intestini* showed very high sensitivity to various pesticides. Its numbers were positively increased for fenhexamid (p), fluopyram (p), clothianidin (n) and metconazole (b). It is worth noting that a positive correlation was found between the pesticides listed. Few species from the *Apilactobacillus* genus showed negative relationships with the fluopyram (n) while some species of the genus *Apibacter* showed a positive relationship with the same pesticide. For boscalid (b), a positive association was found with few opportunistic bacterial organisms: *S. marcessense* and *P. agglomerans*.

As with the ANCOM analysis, the SelBal analysis revealed that *B. intestini* was present in several numerators of the balances, with balances being higher in the presence of pesticides (Supplementary figures 6,7). The balance associated with fluopyram (n) content included a species of *Apilactobacteriа* – *A. sp.wkB309*.

Correlation analysis revealed a wide range of microorganisms associated with various pesticides (Figure 4d). Only for pyroclostrobin (p) was no significant correlation found, as was the case with ANCOM-BC2 and SelBal. For the remaining pesticides, the number of bacteria associated with them was generally higher. Many negative relationships between microbes and pesticides were also found, which may reflect a bias in the method^31,40–42^.

**Figure 4.**
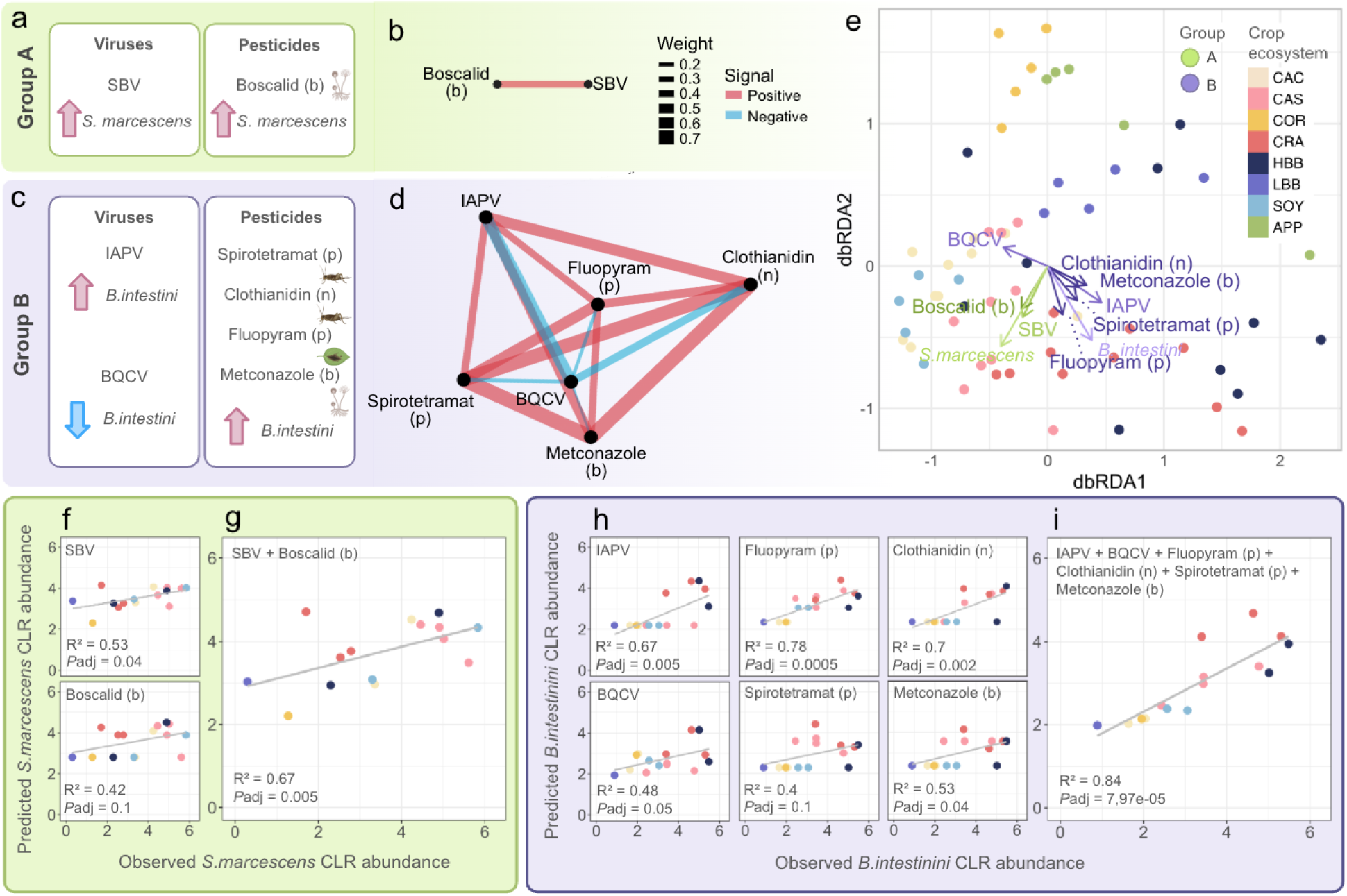
Connection between microbiome, pesticides and viruses (a,c) Schematic summary of significant associations identified for Groups A (a) and B. Group B (c) includes predictors associated with *S. marcescens*: SBV and boscalid (b). Group B includes predictors associated with *B. intestini*: IAPV, spirotetramat (p), clothianidin (n), fluopyram (p), and BQCV with negative association. Arrows indicate direction of association (increase or decrease in CLR-transformed abundance). **(b,d)** Correlation network for Group A and Group B predictors. Edge color indicates direction of association (red – positive, blue – negative), and edge thickness reflects correlation strength. **(e)** Distance-based redundancy analysis (db-RDA) ordination plot. Arrows indicate significant predictors shaping microbiome variation. Samples are colored by the crop ecosystem and grouped according to metadata. Vector direction reflects increasing predictor values; vector length indicates strength of association. **(f-g)** Predictive performance of the linear regression model for *S. marcescens* based on the features from the Group A. The transformed CLR count of bacteria was used as a predictor. The graphs differ in the samples supplied to the training and test datasets. **(h-i)** Predictive performance of the linear regression model for *B. intestini* based on the features from the Group B. The transformed CLR count of bacteria was used as a predictor. The graphs differ in the samples supplied to the training and test datasets. **(f-i)** 75% of the samples from the original dataset were used as the training dataset. The remaining 25% of the samples were used as the test dataset. Prediction of transformed bacterial abundance based on the number of viruses and pesticides separately **(f,h)** and merged **(g,i)**

When combining the results, several bacterial organisms were identified that showed similar patterns of association with pesticides across different analysis types (Table 2, Figure 3b). For example, *B. mellis* showed a negative association with fenhexamid-contaminated pollen in all three analyses. *B. intestini*, in contrast, was positively associated with three pesticides: fluopyram (p), spirotetramat (p), and clothianidin (n). Only one bacterium, *A.* sp. wkB309, was associated with fluopyram (n) across all analyses, and was present in higher abundance when this pesticide was used. For boscalid (b), the opportunist *S. marcescens* and the core organism *Gilliamella strain A7* were present in higher numbers.

**Table 2.** Common microorganisms found in different analysis types. The rows show the combinations of analyses. The columns indicate viruses for which positive results were found if the pesticide and bacterial counts increased simultaneously, or negative results if the bacterial counts increased in the absence of the pesticide. In the case of SelBal, a microorganism was classified as positively associated if it was present in the “numerator” group and negatively associated if it was in the “denominator” group.

| Pesticide | Fenhexamid pollen |  | Fluopyram pollen |  | Spirotetramat pollen |  | Difenoconazole pollen |  | Pyraclostrobin pollen |  |
| --- | --- | --- | --- | --- | --- | --- | --- | --- | --- | --- |
| Effect upon exposure | Positive | Negative | Positive | Negative | Positive | Negative | Positive | Negative | Positive | Negative |
| ANCOM-BC2 x | *S. melliferum* | - | *B. intestini* | - | *B. intestini* | - | - | - | - | - |
| SelBal |  |  |  |  |  |  |  |  |  |  |
| ANCOM-BC2 x Correlations (clr) | - | - | *B. intestini* | - | *P. alvei*<br>*B. intestini* | - | - | *G. apis* | - | - |
| Correlations (clr) x SelBal | - | *B. mellis* | *B. intestini* | *S. apis* | *B. intestini* | *B. actinocoloniiforme* | *A. apis*<br>*B. intestini* | - | - | - |
| **General species** | - |  | *B. intestini* | - | *B. intestini* | - | - | - | - | - |
| General counts | 0 | 0 | 1 | 0 | 1 | 0 | 0 | 0 | 0 | 0 |

| Pesticide | Fluopyram nectar |  | Clothianidin nectar |  | Metconazole bees |  | Boscalid bees |  |
| --- | --- | --- | --- | --- | --- | --- | --- | --- |
| Effect upon exposure | Positive | Negative | Positive | Negative | Positive | Negative | Positive | Negative |
| ANCOM-B C2 x SelBal | *A. sp.wkB309* | - | *B. intestini* | - | *B. intestini* | - | *S. marcescens*<br>*G. sp.A7* | - |
| ANCOM-B C2 x Correlations (clr) | *A. sp.wkB309*<br>*A. adventoris*<br>*A. mensalis* | - | *B. intestini* | - | *B. intestini* | - | *S. marcescens*<br>*P. agglomerans*<br>*G. sp.A7* | - |
| Correlations (clr) x SelBal | *G. sp.A7*<br>*A. sp.wkB309* | *L. helsingborgensis* | *B. intestini* | *B. actinocoloniiforme* | *B. intestini* | - | *S. marcescens*<br>*G. sp.A7* | *L. helsingborgensis*<br>*L. melliventris*<br>*L. sp.ESL0261* |
| **General species** | *A. sp.wkB309* | - | *B. intestini* | - | *B. intestini* | - | *S. marcescens*<br>*G. sp.A7* | - |
| General counts | 1 | 0 | 1 | 0 | 1 | 0 | 2 | 0 |

For difenoconazole (p), pyraclotrobin (p), and metconazole (b), no bacteria were found that were associated with these pesticides.

Overall, the results show that 6 pesticides were associated with 5 bacterial organisms. Moreover, one of the organisms was associated with 3 pesticides, while in another case, one pesticide affected 2 bacterial organisms.

### Combined effects of viruses and pesticides on microbial community structure

As a result of our analysis, we observed patterns of overlapping effects: some bacteria were associated with multiple factors, and certain factors influenced more than one taxon. It is seen that *S. marcescens* and *B. intestini* showed associations with multiple features. Those features were grouped into clusters A and B (Figure 4a).

For factors within the created groups, a correlation analysis was conducted to find the relationship between them. Within group A, a positive correlation was detected between boscalid (b) and SBV (r = 0.4; <0.05) (Figure 4b). In group B, clothianidin (n) and spirotetramat (p) were strongly correlated (R² = 0.7; p <0.05) and jointly associated with IAPV (R² = 0.6; p < 0.05) (Figure 4c). Fluopyram showed a moderate association with these parameters (R² = 0.4; p < 0.05). In contrast, BQCV exhibited negative correlations with the listed factors and was more abundant in the absence of clothianidin (n) and under low IAPV infection levels.

Also, to reflect these parameters in relation to each other and microbes, as well as reflecting their general relationship with microbial communities, we conducted an db-RDA analysis. Distance-based redundancy analysis (db-RDA) confirmed the coordinated influence of these predictors on microbiome structure. In group A, the vectors corresponding to SBV (Variance = 4.079; F = 3.52; p = 0.002) and boscalid (b) (Variance = 3.354; F = 2.90; p = 0.007) were aligned, as was the vector for *S. marcescens* (Figure 4d). In group B, vectors for IAPV, spirotetramat (p), fluopyram (p) and clothianidin (n) pointed in a similar direction and coincided with *B. intestini*, whereas BQCV was oriented oppositely (Figure 4d). Among all predictors, BQCV (Variance = 5.131; F = 4.43; p = 0.001) and IAPV (Variance = 3.854; F = 3.33; p = 0.004) explained the largest proportion of variance, but spirotetramat (p), fluopyram (p) and clothianidin (n) did not show significant associations with microbiome variation (0.5 < p < 0.8).

Given the identified relationships between viruses, pesticides, and bacterial taxa, we hypothesized that, taken together, they should have stronger predictive power than separately on the current dataset. For the group A predictors, a linear regression model was trained using a 75/25 split of training and test sets. The model was trained using the boscalid (b) and SBV virus parameters separately (Figure 4f) and together (Figures 4g) as features and *S. marcessens* with CLR-transformed abundance as a predictor. After training the model on the training set (45 samples), we applied the test set. We then compared the predicted values for Serratia on the test set with the actual values and obtained a correlation coefficient of r² = 0.67 and *p* = 0.005. In contrast to the test on both factors, we observed that the model trained separately on viruses and pesticides yielded lower predictive ability (r² = 0.53, p = 0.04 and r² = 0.42, p = 0.1, respectively). Despite the accuracy of the obtained results, in both cases we observed a fairly high RMSE (1.1067) for the observed SD (1.5).

The same analysis was constructed for group B predictors. For combined features it showed explanatory power R² = 0.84, p=0.01 (Figure 4i). For individual stressors, the predictive ability was on average equal to R² = 0.59 with p<0.05 (Figure 4h). As a result, similar to the results obtained for Group A, we found that when using pesticides and pathogens separately, the predictive model performed worse than when using the factors together. In both cases, the model also had a fairly high RMSE at the existing SD.

For both groups our analysis revealed some clustering by the crop ecosystem. This is logical, as we initially assumed that the crop ecosystem has a significant relationship with fungi and bacteria, but we need to demonstrate that this relationship will persist. To assess the robustness of the identified associations and test their replicability we have used external data.

### Validation results on multiple external datasets

To assess the reproducibility of the identified relationships, we predicted bacterial abundances on two additional datasets with different spatial and temporal coverage (Figure 5a)^34^. We then assessed the correlation between predicted and expected values, similar to the testing in the previous section. This approach allowed us to determine which relationships between the studied factors and bacterial taxa persist beyond the original sample and can be considered the most reliable.

**Figure 5.**
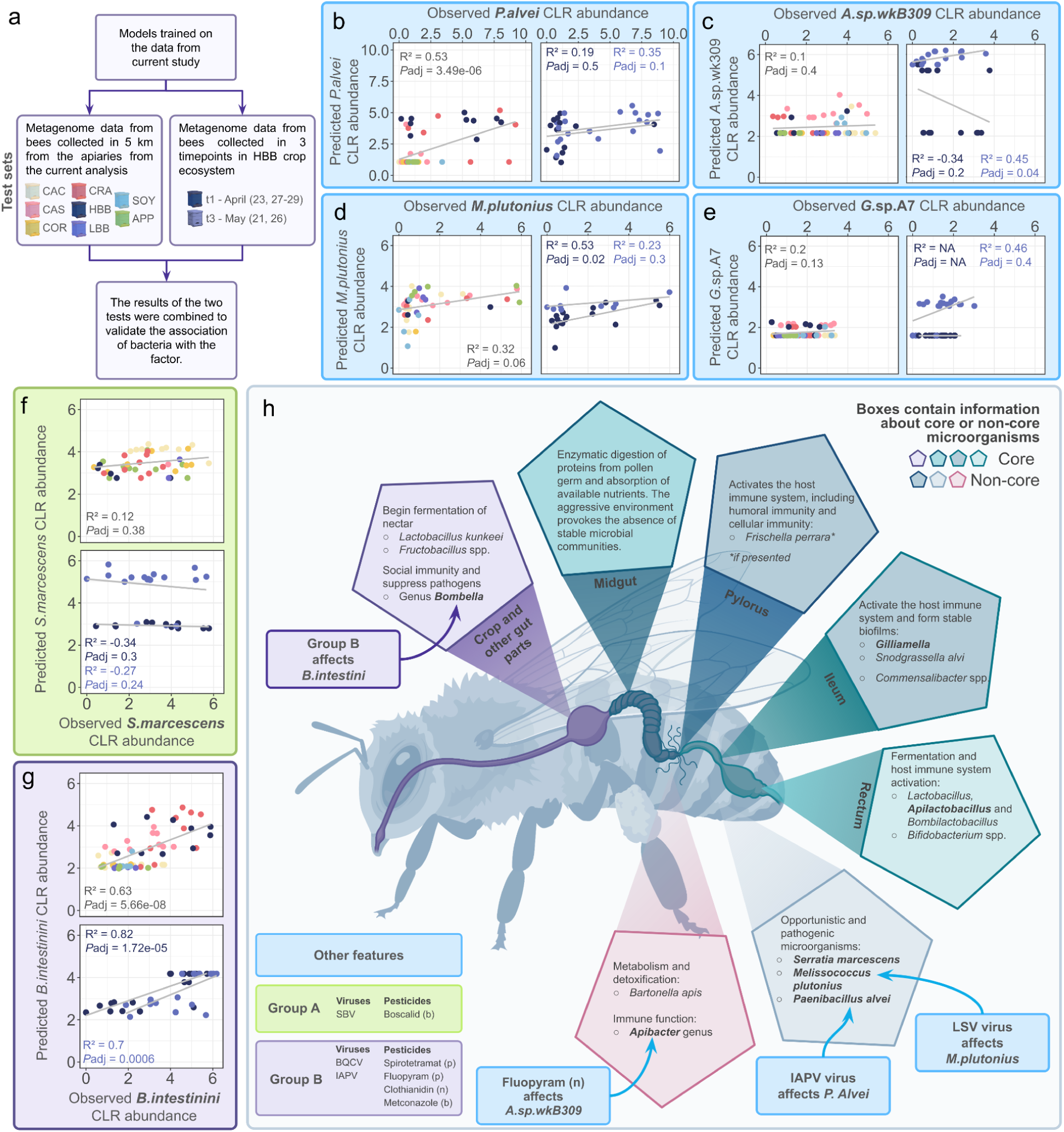
Reproducibility of bacterial associations across independent metagenomic datasets. **(a)** Validation scheme for bacterial associations with the studied factor. Models trained on data from the current study were tested on two independent metagenomic datasets: a sample of bees collected within a 5-km radius of the studied apiaries and a sample from the HBB agroecosystem collected at three time points. The results from both tests were combined to confirm the association of bacteria with the factor. **(b-g)** Validation of model predictions on independent datasets for individual bacterial taxa. The graphs show the relationship between the observed and predicted CLR abundances of *P. alvei* **(b)**, *A.* sp.wkB309 **(c)**, *M. plutonius* **(d)**, *G.* sp.A7 **(e)**, *S. marcescens* **(f)** and *B. intestini* **(g)**. For each taxon, the results of model testing on independent samples are presented, including the determination coefficient (R²) and the adjusted significance value (*Padj*). The resulting relationships reflect the degree of reproducibility of the model predictions on external data and are used to confirm the association of the corresponding bacteria with the factor under study. **(h)** Conceptual model summarizing the effects of viruses and pesticides on gut bacterial taxa in bees. Boxes indicate core and non-core microorganisms and their functional roles within gut compartments. Arrows illustrate the direction of influence of specific stressors on bacterial taxa, highlighting predominant effects on opportunistic microorganisms.

For *P. alvei*, partial confirmation of the results is observed. In a test on bees sampled far from the apiaries, the association was quite pronounced and statistically significant (R² = 0.57, Padj = 3.49e-06), although a significant number of false negative zero values were observed (Figure 5b). However, in the subsamples of time points, the association was reproduced close to statistical significance only at t3. For *A.* sp.wkB309, confirmation was also incomplete (Figure 5c). Only for the t3 subset of time points was statistically significant prediction accuracy demonstrated (R² = 0.45, Padj = 0.04), but in the remaining subsamples, the results were weak or uninterpretable. However, if only those samples without zero pesticide values were included in the training and test datasets, the prediction would be statistically accurate. For *M. plutonius*, a more confident external validation is observed: in the test on samples removed from the hives and on samples taken in the t1 time interval, positive and statistically significant relationships were noted (R² = 0.32, Padj = 0.04; R² = 0.53, Padj = 0.02, respectively) (Figure 5d). For *G.* sp.A7, the results were not reproducible (Figure 5e). In some tests, correlations were absent or could not be reliably estimated, and statistically significant confirmation was not obtained.

For *S. marcescens* validation using external data did not confirm the initial association: R² values were low or negative, and Padj was insignificant (Figure 5f). Therefore, for this taxon, the results were not replicated in independent datasets, and we cannot be certain of their relationship. However, *B. intestini* showed the most convincing reproducibility. In 3 of 4 tests, high levels of prediction correlation and low levels of *p* were observed (R² = 0.63, Padj = 5.66e-08; R² = 0.82, Padj = 1.72e-05; R² = 0.70, Padj = 0.0006) (Figure 5g). This allows us to consider *B. intestini* as one of the most reliably supported taxa in this analysis, although we observe that confirmation in t4 was not found.

Based on these findings, we developed a schematic representation illustrating the effects of viruses and pesticides on bacterial taxa, accounting for their localization within the bee gut (Figure 5h). In most cases, the identified factors primarily affected opportunistic microorganisms, including *Serratia* and *Melissococcus*. Members of the core microbiota were substantially less sensitive to pesticide exposure or viral infection and were only occasionally associated with these stressors.

## Discussion

Here, we provide the first complex description of symbiont balances in the gut of honey bees and their maintenance under diverse environmental conditions of Canadian agroecosystems. We show that bee symbionts have statistically significant differences in microbiota abundances in response to factors such as presence of bee viruses and pesticide exposure.

Our samples show the canonical adult honey-bee gut community with the expected low-diversity high-stability core species. In many individuals we also detected typical, lower-abundance taxa such as *Bartonella apis*, *Frishella perrara*, and *Commensalibacter sp.*, all within the ranges reported for healthy workers during the pollinating season^21^. This overall compositional stability provided a baseline against which the influence of stressors could be evaluated.

Preliminary analysis of those stressors showed a strong association between pesticide levels and the crop ecosystem. These patterns may reflect regional differences in pathogen pressure, as some crop ecosystem sampling areas may be located in regions with higher infection prevalence, or they may result from crop-specific pesticide application regimes and seasonal management practices^44,45^. As with pesticides, pathogen analysis revealed that the abundance of bee viruses varied across crop ecosystems. These findings indicate that viral dynamics are not uniform even among geographically proximate or related crop systems. This pattern can likely be explained either by the seasonality of viral loads, or beekeeping management, or hygienic stock or by the different prevalence of viruses across regions, as some bees simply had these viruses initially, while others did not. Overall, the observed variation in viral load reflects the combined influence of crop ecosystem, geographic region, and additional environmental factors, consistent with patterns previously reported by McAfee *et al.* (2025)^43^.

Because we found a strong link between stressor parameters and the crop ecosystem, we conducted analyses in several different ways: without adjusting for the crop ecosystem, with adjusting for the crop ecosystem, and individually for several crops to confirm the observed relationship without adjusting for this parameter. In all analysis options, we observed similar trends in the association of the microbiome with pesticide and pathogen parameters. Thus, pesticides and pathogens connected to bacterial organisms were identified and some of them can be grouped because they affect one common bacterium.

In this way, one of the groups was significantly associated with *Serratia marcescens* (Group A), and the other with *Bombella intestini* (Group B). The existence of such groups is explained by the fact that microorganisms are exposed to multiple factors simultaneously. Group A contains SBV and the fungicide boscalid (b), since both were associated with increasing opportunistic *S. marcescens*. This pattern indicates a simultaneous increase of bacterial and viral infection with boscalid exposure. Boscalid may increase virus load, which could trigger a secondary bacterial infection. Alternatively, boscalid may reduce bee immunity and increase the abundance of opportunistic pathogens including *Serratia marcescens*, which may in turn lead to an increase in viral load. It is important to note that other studies have reported varied responses to boscalid by the microbial community of the intestine and cuticle of bees^30,32,33,46,47^.

Group B included IAPV and BQCV, linked to the pesticides fluopyram (p), spirotetromat (p) and clothianoidin (n) and metconazole (b). Those stressors, except BQCV, increased *B. intestini* abundance. This shows a possible depletion of beneficial gut symbionts under BQCV infection. As *B. intestini* was positively associated with multiple pesticides and IAPV, it indicates either relative tolerance of this species or indirect competitive release under pesticide exposure. This finding aligns with prior observations that *Bombella* spp. often increase in disturbed or chemically stressed gut environments, potentially reflecting metabolic flexibility or resistance mechanisms^48,49^. In a previous multi-omic analysis that included only highbush blueberry (HBB) and cranberry (CRA) agroecosystems we also found that clothianidin detection in bees was associated with an increase of *B. intestini*, as well as *Melissococcus plutonius*^33^. Further, in a separate study where we experimentally exposed bees to clothianidin, we also found a significant increase of *Bombella apis* (previously *B.* sp.ESL0368^50^) as well as *Paenibacillus alvei*^29^. These results indicate that different *Bombella* species may respond to clothianidin and other pesticides depending on exposure conditions or the presence of additional concurrent stressors, such as LSV or IAPV viruses. In the current study, LSV virus was corepresented with *Melissococcus plutonius*, an association previously reported by Hesketh-Best *et al.* (2024)^28^. High levels of IAPV virus also caused an increase of the opportunistic bacterium *Paenibacillus alvei*, suggesting a robust relationship between LSV and IAPV and shifts in both core and opportunistic gut bacteria, particularly those associated with the development of European foulbrood disease^51^. Pesticides were also found to affect the microbiota. In particular, *Bartonella mellis* showed a reproducible negative association with fenhexamid detected in pollen, suggesting sensitivity of this gut symbiont to fungicide exposure. Previous studies have similarly reported declines in *Bartonella* spp. following exposure to agricultural fungicides and insecticides, consistent with their known susceptibility to xenobiotics and their reliance on host-derived carbohydrates for persistence in the gut ecosystem^52^. Interestingly, fluopyram, detected in nectar, had different connected bacteria, than fluopyram found on the pollen, and was connected to the *Apibacter* sp. wkB309. Fluopyram detection in nectar and pollen did not always co-occur in samples from the same site, which may explain the variable effects on different bacterial organisms, particularly those with differences in localisation. Together, these findings highlight that different viruses and pesticides are associated with distinct microbial signatures rather than a uniform dysbiosis response. The enrichment of opportunistic or pathogenic bacteria in the presence of certain viruses may indicate that viral infection creates ecological niches that favor bacterial dysbiosis, potentially exacerbating disease severity. Conversely, changes involving microbiota members may affect viral infections by interfering with beneficial host–microbe interactions that are important for immune function and gut homeostasis.

It is important to note that the reproducibility of the results was not confirmed for all bacterial taxa. This discrepancy may be due to both the biological context-dependence of individual associations and differences between independent samples in environmental conditions, sampling time points, community composition, and sequencing depth. Taken together, this suggests that some associations may be robust and biologically significant, while others require further validation using larger, more standardized datasets.

### Limitations of the study

It is important to note that our research used the BeeRoLamMa database with 55 distinct taxa based on the known bee gut microbiome composition at the time the database was created. Even though the taxa list has since increased as reflected in the European Nucleotide Archive (ENA) database, most reads were assigned to the honey bee host genome or to the microbial taxa listed in the database used with few reads not related to the bee or bacterial community.

## Materials and methods

### Sample collection

Samples for metagenomic sequencing were collected from sites across Canada that represented the country’s diverse climate and landscape as described in Tran et al. (2025)^34^. The analysis used samples collected from colonies that were placed near (<0.5 km) focal crops: apples (Quebec, 2021), seed canola (Alberta, 2020 and 2021), commodity canola (Manitoba, 2020 and 2021, and Alberta, 2020 and 2021), corn (Ontario, 2020) cranberries (British Columbia, 2020, and Quebec, 2021), highbush blueberries (British Columbia, 2020 and 2021) (Fig 1.A), lowbush blueberries (Quebec, 2021), and soybeans (Manitoba, 2020). Sampling took place during the crop bloom, the timing of which varied for each crop in the ’mid-bloom’ for the crop. At each sampling site during bloom, nurse-aged bees were collected from each of four colonies on site, transferred to dry ice, and stored at -80 °C until used. The intestinal tract of twelve bees per site - three randomly selected nurse bees from each of the 4 colonies in a site - was removed, pooled, and homogenized, resulting in 60 samples for the multi-crop experiment as described in Tran et al. (2025)^34^.

### Microorganisms classification and abundance estimation

For taxonomic classification, raw sequencing reads were adapter-trimmed and quality-filtered using Fastp (v0.23.2). Reads were taxonomically classified as bacterial using Kraken 2 (v2.1.2) 56 in conjunction with the BeeRoLaMa v1 Honey Bee Microbiota Database^34,53^.

For *Lactobacillus* genera we rebuilt the taxonomic classification according to the new 23 *Lactobacillus* genera article^54^. Species strains *C.* ESL0284 and *C.* ESL0366 were merged into two *C. melissae* based on NCBI classification.

### Determination of the amount of pesticides and pathogens

From each colony, another batch of worker bees (n = 12/colony) were sampled and pooled for pathogen and agrochemical residue analysis. Pathogen detection followed OIE and COLOSS Bee Book guidelines. Major honey bee viruses assessed via real-time qPCR, primers and detailed methods are available in a previous report in the Supplementary Data 2 of McAfee *et al.* (2025)^43^. From pathogens for further analysis only highly representative variables were taken, which were detected in more than 30% samples (black queen cell virus = BQCV, deformed wing virus types A and B = DWV-A and DWV-B, Israeli acute paralysis virus = IAPV, sacbrood virus = SBV and Lake Sinai virus = LSV). Other variables were assumed to be noise and removed from the analysis. All virus pathogens variables were normalised with log(x+1) to bring the data closer to a normal distribution. After normalisation we assessed the normality of pathogenic variables using the Shapiro-Wilk Test from the stat package as the sample size is n = 60. The package ggplot^55^ was used to visualize pathogen distribution for each viral pathogen.

To examine the association between viral pathogens and the honey bee gut microbiome, we analyzed existing viral infection data. Nine viruses were initially considered, but to ensure sufficient representation, only those detected in at least 30% of samples were included in downstream analysis.

Additionally, agrochemical residue analysis using LC-MS/MS at the Agriculture and Food Laboratory, University of Guelph, an ISO/IEC 17025 accredited laboratory, was performed. The analysis covered 232 unique pesticide compounds. Strict quality control measures, including method blanks, blind proficiency testing, and second-person validation, were implemented to ensure data accuracy and to prevent transcription errors. Similar to our viral analysis, only highly representative pesticide variables detected in more than 30% samples were analyzed further.

We investigated the relationship between the microbiome and pesticides using all available exposure measurements. As with pathogens, of the 132 pesticides initially considered, we limited the analysis to compounds detected in at least 30% of samples to ensure stable estimates and reduce noise from null detections^56,57^. Eight pesticides met this abundance criterion: 5 were detected in pollen, 2 in nectar, and 1in bee tissue.

### Analysis of microbiome in different crop ecosystems

Analyses of microbial community composition, alpha- and beta-diversity metrics and statistics were performed using R (version 4.4.2). Relative abundance was calculated and genera which had an average proportion of less than 0.01% of counts were not included in further analysis. Rarefaction, diversity analysis of microbial community composition were performed with the vegan and base R packages^58^.

Relative abundances were used to describe microorganism composition. For visualisation, microbial taxa were grouped at the genus level. Plots were generated using the ggplot2 barplot packagefor the relative abundance data and pheatmap package (version 1.0.1^59^)for centered log ratio transformed (clr)33 data. Microbiome transformation was performed with the transform function from the microbiome package (microbiome::transform(data, “clr”^60^). Sample and genus clustering in the heatmaps utilised Euclidean distances on log10-transformed data (cluster_rows and cluster_cols parameters).

Beta-diversity distances and Principal Coordinate Analysis (PCoA) visualisations were calculated using the dist function from the stats package (version 4.4.2). Multivariate homogeneity of group variances was assessed using the betadisper function, and partitioning of distance matrices among sources of variation with linear model fitting was performed using the adonis2 function (vegan package version 2.7^58^; see part ‘Beta-diversity’ in “BeeCsiMetagenome” on GitHub https://github.com/andwhoami/BeeCsiMetagenome) (Dixon, 2003). Classical multidimensional scaling (MDS) for PCoA visualisation was performed using the cmdscale function (stats package). Visualisations were generated with the geom_point function from the ggplot2 package (version 3.5.2)^55^. Alpha diversity indices (Simpson, Shannon, Chao1) were computed using the vegan package^58^. Comparisons were performed using the Wilcoxon Test, with *p* values adjusted using the Benjamini-Hochberg method.

To assess the cause of differences in beta-diversity, differential abundance analysis of transformed bacteria was performed using central relative logarithm and assessed using the Wilcoxon-Test and Student’s t-test and ANCOM-BC2 analysis^36^. To confirm the significance of the differences, we sampled the microorganisms included in the analysis on untransformed data. The resulting samples were transformed by clr and the comparison of the averages was repeated also by the Wilcoxon-Test and Student’s t-test.

### Explanation of pathogens and pesticides connection to the different сrop ecosystems

To test associations between the bee crop community and virus and pesticides loads/presence, we quantified community beta-diversity from rarefaction-free abundance tables using both Bray–Curtis (composition/abundance-sensitive) and Jaccard (presence–absence) distances, visualized by PCoA. We then evaluated virus effects on crop composition with PERMANOVA (999 permutations) on each distance matrix for analysis of variance while controlling for covariates and stratifying permutations by colony where appropriate. To avoid inflated effects from correlated predictors, we assessed collinearity among viral and environmental variables using variance inflation factors (VIF) and pairwise correlations, iteratively removing or combining variables until all VIF < 3. Eta-squared (η²) was computed directly from the ANOVA sums of squares as the proportion of total variance attributable to a factor.

We quantified how much of the microbiome variation was explained by crop composition and viruses separately and jointly using variance partitioning with partial distance-based RDA (db-RDA). From the same Aitchison distance matrices, we fit three models: (a) crop only (conditioning on covariates), (b) viruses only (conditioning on covariates), and (c) crop + viruses together. Also we quantified crop–virus overlap in explaining community composition using the betadisper and adonis2 functions on the same distance matrix. We also assessed the correlation of factors with themselves (viruses with viruses, pesticides with pesticides) using the Spearman’s rank correlation.

Given that we found a strong link between environmental parameters and the crop ecosystem, we conducted subsequent analyses in several different ways: without adjusting for the crop ecosystem, with adjusting for the crop ecosystem, and individually for several crops to confirm the observed relationship without adjusting for this parameter.

### Explanation of microorganisms-metadata connection

We used a few methods and their comparison to explain the variation of microbiome and its connection to the metadata. To assess the relationship of microbes with viruses and pesticides, we used the ANCOM-BC2 analysis, Selbal analysis, and correlation analysis on the CLR transformed data^36,40^.

We identified differentially abundant taxa with ANCOM-BC2 (R package ancombc2) using the raw count table (no rarefaction) and the package’s bias-corrected log-linear model. Initially, we included only uncorrelated viral pathogens in the model and did not adjust for the crop ecosystem on the uncentered data. For visualisation we used only significant bacterial organisms. We then took the centered data and adjusted for the crop ecosystem. We also conducted a similar analysis for HBB in the CRA separately to demonstrate that the observed differences persisted.

We analyzed compositional microbiome data with SelBal, which identifies an interpretable log-ratio (“balance”) between two sets of taxa that best associates with an outcome. After zero replacement to enable logs, SelBal performs greedy forward selection: it first evaluates all single-taxon pairs to form an initial balance, then iteratively adds one taxon at a time to either the numerator or denominator, choosing the addition that maximally improves predictive/associative performance, while optionally adjusting for covariates. We used SelBal on the data on the presence or absence of pesticides or pathogens and did for each variable.

For the correlation analysis, we used all the data. We calculated the correlation coefficients and the *p* values for them and visualized only the statistically significant ones. We used Pearson and Spearman rank correlations on log-ratio transform (CLR) data with logarithmised metadata variables. As far as CLR transforms compositional data into real Euclidean space, it enables the use of classical tools like correlation, regression, and clustering. For *p* values adjusted Benjamini-Hochberg control was used. To visualise correlation we used Pearson rank correlation results with *P _adj_* values. We visualise microbiome CLR transformed data with log10 pesticides and pathogens separately using heatmap function with star *p* values.

To run and average the selected analysis methods, we constructed a R library. The library included the SelBal and ANCOM-BC functions, as well as the calculation of correlations on the CLR of transformed data. The main function aggregates all results and creates an object indicating the bacteria common to each analysis and the side of influence of each.

### Explanation of microbiome connection to pesticides and viruses

Pesticides and viruses, connected to the same bacterial organisms, were grouped into two clusters (A and B) corresponding to taxa influenced by multiple factors. For features inside the groups correlation analysis was performed to investigate connections between those parameters. Visualisation was performed with ggraph from ggplot2^55^. To verify the stability of detected correlations, analyses were repeated using a geographically restricted subset of samples.

To evaluate the coordinated effect of predictors on overall microbiome structure, distance-based redundancy analysis (db-RDA) was performed using Aitchison dissimilarity matrices with base R statistics library. Significance of individual predictors was assessed using permutation tests (n = 999 permutations). The proportion of explained variance and F-statistics were calculated for each variable. Ordination plots from the ggplot2 were used to visualize the direction and magnitude of associations between predictors and bacterial taxa.

To assess predictive capacity, linear regression models were constructed using CLR-transformed bacterial abundances as response variables and merged and separate group-specific predictors as explanatory variables. Data were randomly split into training and test sets (85/15). Model performance was evaluated on the test set using the coefficient of determination (R²) and significance testing. Analyses were repeated using an expanded dataset to assess model robustness and reproducibility.

### Validation results on multiple external datasets

To validate the identified associations, the dataset containing relationships between the studied factors and bacterial taxa was used as a training set from our previous research. For each taxa, a linear model was built using the data from the current study, after which the trained models were applied to two datasets used as test sets^34^. This approach allowed us to assess whether the identified relationships were reproducible beyond the original dataset.

The quality of external validation was assessed based on the degree of agreement between the model’s predicted and observed bacterial abundance values in the test datasets. For this purpose, the correlation between the predicted and actual values was calculated, and the resulting relationships were visualized in scatterplots with linear regression lines plotted in ggplot2^55^. The resulting plots were used to compare the reproducibility of the results between independent samples.

### Schematic representation of the results

The schematic model summarizing virus–pesticide–microbiome interactions was constructed by integrating statistically significant associations identified in this study with published knowledge on bacterial localization and functional roles within the bee gut. Information on spatial distribution of microbial taxa and their functional characteristics was compiled from previously published microbiome and functional studies of *Apis mellifera*^21,23,26,61,62^.

Only taxa with experimentally supported localization or function were annotated in the model. The direction of influence was derived from the statistical analyses conducted here, while functional and anatomical annotations were incorporated from the literature to provide biological context. The schematic therefore represents an integrative conceptual model combining quantitative results from the present analysis with established knowledge of bee gut microbiology.

## Acknowledgments

This work was funded by the Government of Canada through Agriculture and Agri-Food Canada (AAFC) Genomics Research and Development Initiative (GRDI) funding (AAFC J-002368), Genome Canada (LSARP #16420), and the Ontario Genomics Institute (OGI-185). Mass spectrometry infrastructure used here was supported by the Canada Foundation for Innovation, the BC Knowledge Development Fund, the Life Sciences Institute, and Genome BC (374PRO). AKR was supported by the HSE University basic research program. We thank Mikhail Gelfand, Nancy Moran, Fedor Ryabov and Avril Metcalfe-Roach for helpful comments on the manuscript and figures.

## Data availability

The raw metagenome sequences for all T2 samples are available as BioProject PRJNA999720 https://www.ncbi.nlm.nih.gov/bioproject/PRJNA999720/ https://www.ncbi.nlm.nih.gov/bioproject/PRJNA999720/ in the NCBI BioProject database (https://www.ncbi.nlm.nih.gov/bioproject).

## Code availability

https://github.com/andwhoami/BeeCsiMetagenome

## Authors contribution

Conceptualization: AKR, HZ, LJF, AZ, MMG; Methodology: AKR, HZ, LT, LL, MC, JH, TBD, MP, LT, AMR, ASG, AZ, LJF, SFP MMG; Validation: AKR, HZ; Formal Analysis: AKR, HZ, LL, JZ, JH, TBD, MC, RM, HJ, KMM, AJ; Data Curation: HZ, IMC, MP, SKF, LT; Writing – Original Draft: AKR, HZ, LJF, MMG; Writing – Review & Editing: all; Funding Acquisition: LJF, AZ, SFP, MMG; Resources: AZ, LJF, SFP, MMG, ROP, HJ, HZ; Project Administration: AZ, LJF; Supervision: HZ, LJF, AZ, SFP, MMG, HJ, ROP.

## References

1. Khalifa, S. A. M. et al. Overview of Bee Pollination and Its Economic Value for Crop Production. Insects 12, 688 (2021).

2. Discua, S. A. & Longing, S. D. Attractiveness of Drought-tolerant Plants to Insect Pollinators in the Southern High Plains Region. J. Kans. Entomol. Soc. 94, (2022).

3. Rader, R. et al. Non-bee insects are important contributors to global crop pollination. Proc. Natl. Acad. Sci. 113, 146–151 (2016).

4. Stanley, D. A., Msweli, S. M. & Johnson, S. D. Native honeybees as flower visitors and pollinators in wild plant communities in a biodiversity hotspot. Ecosphere 11, e02957 (2020).

5. Maini, S., Medrzycki, P. & Porrini, C. The puzzle of honey bee losses: a brief review. 63(1), 153–160 (2010).

6. Neumann, P. & Carreck, N. L. Honey bee colony losses. J. Apic. Res. 49, 1–6 (2010).

7. Hristov, P., Shumkova, R., Palova, N. & Neov, B. Factors Associated with Honey Bee Colony Losses: A Mini-Review. Vet. Sci. 7, 166 (2020).

8. Gray, A. et al. Loss rates of honey bee colonies during winter 2017/18 in 36 countries participating in the COLOSS survey, including effects of forage sources. J. Apic. Res. 58, 479–485 (2019).

9. Phiri, B. J., Fèvre, D. & Hidano, A. Uptrend in global managed honey bee colonies and production based on a six-decade viewpoint, 1961–2017. Sci. Rep. 12, 21298 (2022).

10. Aizen, M. A. & Harder, L. D. The Global Stock of Domesticated Honey Bees Is Growing Slower Than Agricultural Demand for Pollination. Curr. Biol. 19, 915–918 (2009).

11. vanEngelsdorp, D., Hayes, J., Underwood, R. M. & Pettis, J. A Survey of Honey Bee Colony Losses in the U.S., Fall 2007 to Spring 2008. PLoS ONE 3, e4071 (2008).

12. Espregueira Themudo, G., et al. Declining genetic diversity of European honeybees along the twentieth century. Sci. Rep. 10, 10520 (2020).

13. Glenny, W. et al. Honey bee (Apis mellifera) colony health and pathogen composition in migratory beekeeping operations involved in California almond pollination. PLOS ONE 12, e0182814 (2017).

14. Goodrich, B. K., Williams, J. C. & Goodhue, R. E. The Great Bee Migration: Supply Analysis of Honey Bee Colony Shipments into California for Almond Pollination Services. Am. J. Agric. Econ. 101, 1353–1372 (2019).

15. Sánchez-Bayo, F. et al. Are bee diseases linked to pesticides? — A brief review. Environ. Int. 89–90, 7–11 (2016).

16. Raymann, K. & Moran, N. A. The role of the gut microbiome in health and disease of adult honey bee workers. Curr. Opin. Insect Sci. 26, 97–104 (2018).

17. Hariprasath, K., Mohankumar, S., Sudha, M., Saranya, N. & Saminathan, V. R. The Role of Honeybee Gut and Honey Microbiome in Sustainable Bee and Human Health. J. Pure Appl. Microbiol. 19, 19–33 (2025).

18. Engel, P. et al. The Bee Microbiome: Impact on Bee Health and Model for Evolution and Ecology of Host-Microbe Interactions. mBio 7, e02164–15 (2016).

19. Zheng, H. et al. Metabolism of Toxic Sugars by Strains of the Bee Gut Symbiont Gilliamella apicola. mBio 7, e01326–16 (2016).

20. Emery, O., Schmidt, K. & Engel, P. Immune system stimulation by the gut symbiont *Frischella perrara* in the honey bee (*Apis mellifera*). Mol. Ecol. 26, 2576–2590 (2017).

21. Kešnerová, L. et al. Gut microbiota structure differs between honeybees in winter and summer. ISME J. 14, 801–814 (2020).

22. Motta, E. V. S. & Moran, N. A. The honeybee microbiota and its impact on health and disease. Nat. Rev. Microbiol. 22, 122–137 (2024).

23. Engel, P., Martinson, V. G. & Moran, N. A. Functional diversity within the simple gut microbiota of the honey bee. Proc. Natl. Acad. Sci. 109, 11002–11007 (2012).

24. Motta, E. V. S., Raymann, K. & Moran, N. A. Glyphosate perturbs the gut microbiota of honey bees. Proc. Natl. Acad. Sci. 115, 10305–10310 (2018).

25. Quinn, A. et al. Host-derived organic acids enable gut colonization of the honey bee symbiont Snodgrassella alvi. Nat. Microbiol. 9, 477–489 (2024).

26. Kešnerová, L. et al. Disentangling metabolic functions of bacteria in the honey bee gut. PLOS Biol. 15, e2003467 (2017).

27. Lang, H. et al. Specific Strains of Honeybee Gut *Lactobacillus* Stimulate Host Immune System to Protect against Pathogenic Hafnia alvei. Microbiol. Spectr. 10, e01896–21 (2022).

28. Hesketh-Best, P. J., Fowler, P. D., Odogwu, N. M., Milbrath, M. O. & Schroeder, D. C. Sacbrood viruses and select Lake Sinai virus variants dominated *Apis mellifera* colonies symptomatic for European foulbrood. Microbiol. Spectr. 12, e00656–24 (2024).

29. Tran, L. et al. Neonicotinoid-induced signature dysbiosis identified via metagenomic sequencing of the honey bee gut microbiome. Sci. Rep. 16, 1211 (2025).

30. Hotchkiss, M. Z., Poulain, A. J. & Forrest, J. R. K. Pesticide-induced disturbances of bee gut microbiotas. FEMS Microbiol. Rev. 46, fuab056 (2022).

31. Wizenberg, S. B., et al. Pollen foraging mediates exposure to dichotomous stressor syndromes in honey bees. PNAS Nexus 3, pgae440 (2024).

32. Reiß, F. et al. Fungicides and insecticides can alter the microbial community on the cuticle of honey bees. Front. Microbiol. 14, 1271498 (2023).

33. Zhong, H. et al. Omics Insights Into the Effects of Highbush Blueberry and Cranberry Crop Agroecosystems on Honey Bee Health and Physiology. PROTEOMICS e70033 (2025) doi:10.1002/pmic.70033.

34. Tran, L. et al. Gut microbiome metagenomic sequences of honey bees (*Apis mellifera*) exposed to crops. Microbiol. Resour. Announc. 14, e00731–24 (2025).

35. Gloor, G. B., Macklaim, J. M., Pawlowsky-Glahn, V. & Egozcue, J. J. Microbiome Datasets Are Compositional: And This Is Not Optional. Front. Microbiol. 8, 2224 (2017).

36. Lin, H. & Peddada, S. D. Multigroup analysis of compositions of microbiomes with covariate adjustments and repeated measures. Nat. Methods 21, 83–91 (2024).

37. Ou, Y., Belzer, C., Smidt, H. & De Weerth, C. Methodological recommendations for human microbiota-gut-brain axis research. Microbiome Res. Rep. 3, (2024).

38. P. Vatcheva, K. & Lee, M. Multicollinearity in Regression Analyses Conducted in Epidemiologic Studies. Epidemiol. Open Access 06, (2016).

39. McAfee, A. et al. Higher prevalence of sacbrood virus in *Apis mellifera* (Hymenoptera: Apidae) colonies after pollinating highbush blueberries. J. Econ. Entomol. 117, 1324–1335 (2024).

40. Rivera-Pinto, J. et al. Balances: a New Perspective for Microbiome Analysis. mSystems 3, 10.1128/msystems.00053-18 (2018).

41. French, S. K. et al. Honey bee stressor networks are complex and dependent on crop and region. Curr. Biol. 34, 1893–1903.e3 (2024).

42. Bixby, M. et al. Identifying and modeling the impact of neonicotinoid exposure on honey bee colony profit. J. Econ. Entomol. 117, 2228–2241 (2024).

43. McAfee, A. et al. Regional patterns and climatic predictors of viruses in honey bee (Apis mellifera) colonies over time. Sci. Rep. 15, 286 (2025).

44. Zhang, Y. et al. Pesticide use is affected more by crop species than by crop diversity at the cropping system level. Eur. J. Agron. 159, 127263 (2024).

45. Neugebauer, K. A. et al. Managing fruit rot diseases of Vaccinium corymbosum. Front. Plant Sci. 15, 1428769 (2024).

46. Dong, J. et al. Ternary Mixture of Azoxystrobin, Boscalid and Pyraclostrobin Disrupts the Gut Microbiota and Metabolic Balance of Honeybees (Apis cerana cerana). Int. J. Mol. Sci. 24, 5354 (2023).

47. Paris, L. et al. Honeybee gut microbiota dysbiosis in pesticide/parasite co-exposures is mainly induced by Nosema ceranae. J. Invertebr. Pathol. 172, 107348 (2020).

48. Jin, M. J. et al. Bombella intestini: A probiotic honeybee(Apis mellifera)gut bacterium. J. Insect Physiol. 164, 104836 (2025).

49. Cuesta-Maté, A. et al. Resistance and Vulnerability of Honeybee (Apis mellifera) Gut Bacteria to Commonly Used Pesticides. Front. Microbiol. 12, 717990 (2021).

50. Härer, L., Hilgarth, M. & Ehrmann, M. A. Comparative Genomics of Acetic Acid Bacteria within the Genus Bombella in Light of Beehive Habitat Adaptation. Microorganisms 10, 1058 (2022).

51. Pannullo, A., Muturi, E. J. & Dunlap, C. A. Characterization of toxin systems of Paenibacillus strains isolated from honeybees. Sci. Rep. 15, 31346 (2025).

52. Rosa-Fontana, A. et al. Bee gut microbiota as an emerging endpoint for the environmental risk assessment of pesticides. Sci. Total Environ. 993, 179977 (2025).

53. Wood, D. E., Lu, J. & Langmead, B. Improved metagenomic analysis with Kraken 2. Genome Biol. 20, 257 (2019).

54. Zheng, J. et al. A taxonomic note on the genus Lactobacillus: Description of 23 novel genera, emended description of the genus Lactobacillus Beijerinck 1901, and union of Lactobacillaceae and Leuconostocaceae. Int. J. Syst. Evol. Microbiol. 70, 2782–2858 (2020).

55. Wilkinson, L. ggplot2: Elegant Graphics for Data Analysis by WICKHAM, H. Biometrics 67, 678–679 (2011).

56. Mesnage, R. et al. Impacts of dietary exposure to pesticides on faecal microbiome metabolism in adult twins. Environ. Health 21, 46 (2022).

57. Xiong, S. et al. Prenatal exposure to trace elements impacts mother-infant gut microbiome, metabolome and resistome during the first year of life. Nat. Commun. 16, 5186 (2025).

58. Dixon, P. VEGAN, a package of R functions for community ecology. J. Veg. Sci. 14, 927–930 (2003).

59. Kolde, R. pheatmap: Pretty Heatmaps. 1.0.13 10.32614/CRAN.package.pheatmap (2010).

60. Lahni, L. microbiome R package.

61. Kwong, W. K. & Moran, N. A. Gut microbial communities of social bees. Nat. Rev. Microbiol. 14, 374–384 (2016).

62. Yun, J.-H., Lee, J.-Y., Hyun, D.-W., Jung, M.-J. & Bae, J.-W. Bombella apis sp. nov., an acetic acid bacterium isolated from the midgut of a honey bee. Int. J. Syst. Evol. Microbiol. 67, 2184–2188 (2017).

