## Supplementary information for "Honey bee (*Apis mellifera*) gut microbiome associations with viruses and pesticides across Canadian agroecosystems"

**SUPPLEMENTARY NOTES**	**2**

Supplementary note 1. Differences in bee microbiome across different years	2

Supplementary note 2. Crop ecosystem explanation of the microbiome differences	2

Supplementary note 3. Explanation of crop ecosystem parameter based on HBB and CRA crop ecosystems	4

Supplementary note 4. Pathogen data analysis	5

Supplementary note 5. Ancom-BC2 models for pathogene	7

Supplementary note 6. Pathogene in connection to the microbiome variation	7

Supplementary note 7. Pesticide data analysis	7

Supplementary note 8. Ancom-BC2 models for pesticide	10

Supplementary note 9. Pesticides in connection to the microbiome variation	10

**SUPPLEMENTARY FIGURES**	**12**

### **SUPPLEMENTARY NOTES**

#### Supplementary note 1. Differences in bee microbiome across different years

To analyze the relationship of microbiomes with the sampling year, we selected only those crop ecosystems for which samples were collected in both 2020 and 2021. These crop ecosystems included samples from CAC, CAS, HBB, and CRA (Supplementary note figure 1).We conducted an analysis of homogeneity of variances for each crop ecosystem separately and found no significant difference within-sample variance (betadisper 0.08 < p < 0.43). We also conducted an analysis of differences in variances, and found that crop composition yearly differed significantly only for samples from the CAS group (adonis 0.007 < p < 0.4, ).

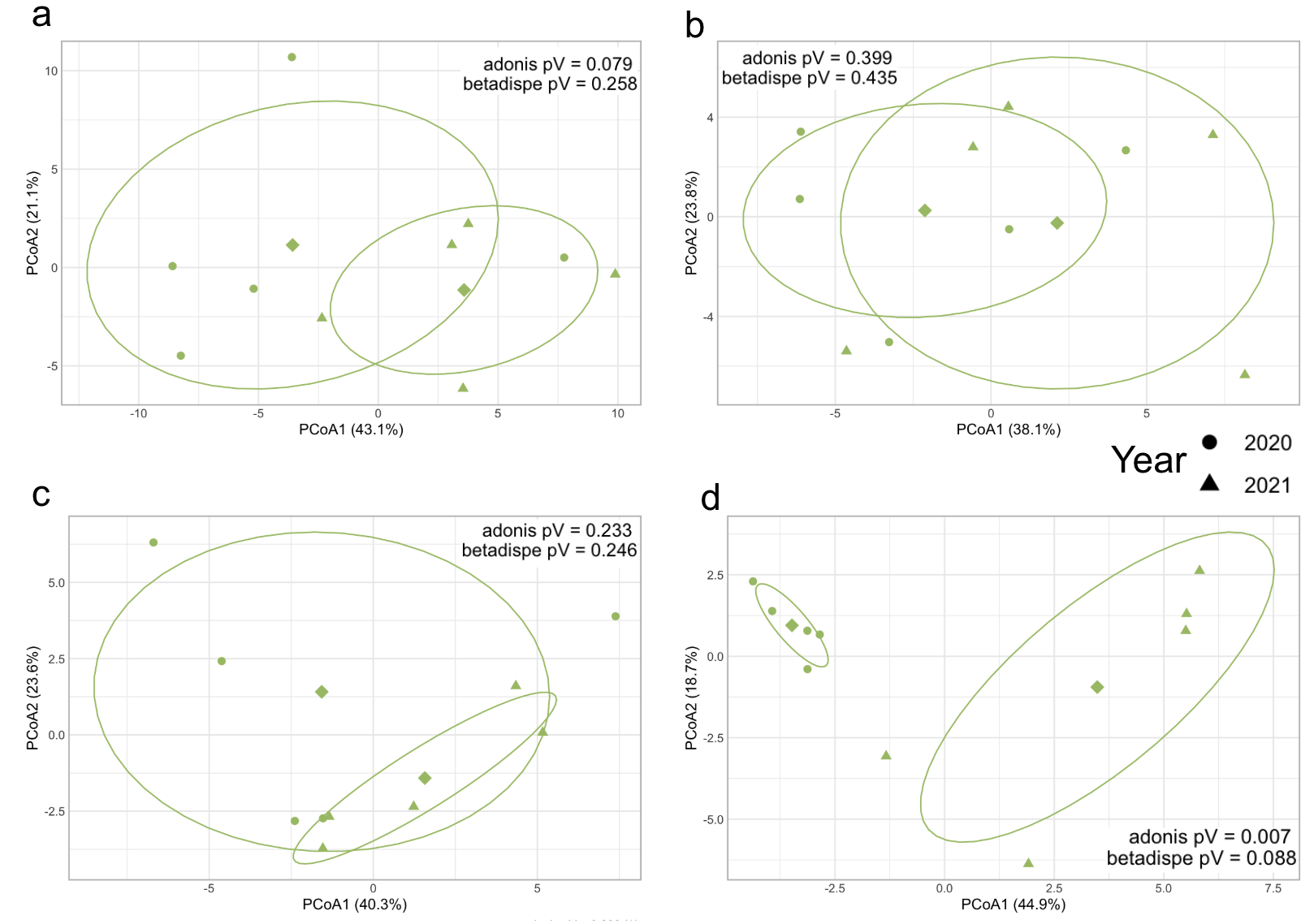

**Supplementary note figure 1.** Plot of the Principal Coordinate Analysis (PCoA) method using the Aitchison distance. The dots are shaped according to the year and panels are connected to different crop ecosystems: a - HBB, b - CRA, c - CAC and d - CAS.

#### Supplementary note 2. Crop ecosystem explanation of the microbiome differences

The differential abundance analysis revealed clear differences in bacterial taxa associated with individual crop ecosystems compared with the pooled group of all other crops (“Other”). Several bacterial species showed strong and consistent enrichment in particular crop systems, while others were depleted relative to the rest of the crop environments (Supplementary note figure 2). Overall, each crop ecosystem exhibited a characteristic set of bacterial taxa with positive log fold change values, indicating higher abundance compared with the combined background of other crops.

In the SOY ecosystem, several taxa were strongly enriched, including *Bartonella apis*, *Pantoea agglomerans*, *Apibacter sp. wkB309*, *Apibacter* sp., and *Serratia marcescens*, all showing positive log fold changes. In contrast, taxa such as *Apilactobacillus kunkeei, Melissococcus plutonius*, and several *Lactobacillus* species were depleted compared with the other crop systems. The APP ecosystem also showed enrichment of multiple taxa, including *Bartonella apis, Pantoea agglomerans*, *Serratia marcescens*, and *Apibacter* species, while *Nosema ceranae* displayed a strong negative log fold change, indicating reduced abundance relative to other crops.

Distinct bacterial signatures were also observed in the remaining crop ecosystems. In the CAC ecosystem, taxa such as *Paenibacillus alvei, Bombella sp.*, and *Lactobacillus* species were enriched, whereas *Bartonella apis* and *Apibacter* sp. wkB309 were strongly depleted. The LBB and HBB crop systems both showed pronounced enrichment of *Bartonella apis*, with additional enrichment of *Spiroplasma melliferum*, *Paenibacillus alvei*, and *Gilliamella* species. In contrast, the clover/rape-associated system CRA showed overall depletion of the examined taxa relative to other crops. The CAS ecosystem displayed moderate enrichment of several *Lactobacillus* and *Apilactobacillus* taxa, while the COR ecosystem showed broad enrichment across many taxa, including *Melissococcus plutonius, Bombella intestini*, *Apilactobacillus kunkeei*, and *Pantoea agglomerans*. These patterns indicate that each crop ecosystem supports a distinct microbial community structure, with specific bacterial taxa preferentially associated with particular crop environments.

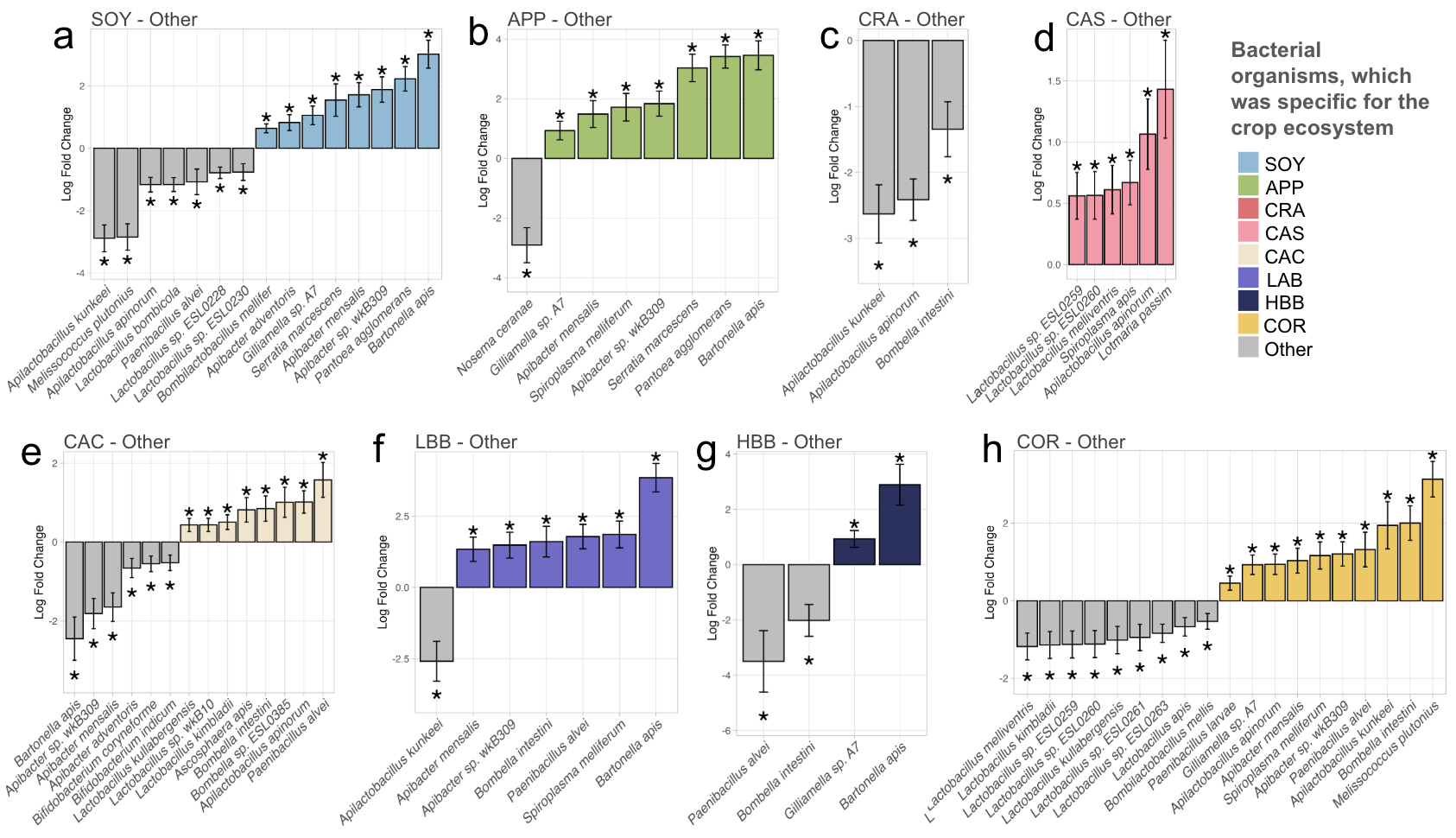

**Supplementary note figure 2.** Log-fold change (LFC) of species abundance in the different crop ecosystems across all samples and years. Statistical significance (ANCOM-BC2; p <0.05) is indicated by *. Positive LFC indicates an increase in abundance in the crop ecosystem of interest.

#### Supplementary note 3. Explanation of crop ecosystem parameter based on HBB and CRA crop ecosystems

To demonstrate that a productive ecosystem is actually a combination of various factors, we analyzed the abundance of crops in two ecosystems. We plotted the abundance of highbush blueberries and cranberries in the ecosystems corresponding to these crops. We found that highbush blueberries were present in both the corresponding ecosystem (HBB) and the productive ecosystem with cranberries (CRA) (Supplementary note figures 3a-b). Moreover, we found that the abundance of highbush blueberries in the productive ecosystem with cranberries was not always lower than in the productive ecosystem with blueberries (HBB) and based on statistics was equal in both crop ecosystems (p=0.16). Cranberries, on the other hand, were present only in the productive ecosystem with cranberries (CRA).

We also analyzed the abundance of certain pesticides and pathogens for this analysis to demonstrate that they differ depending on the productive ecosystems of CRA and HBB. It is clear that there is a statistically significant difference between the abundance of pesticides and pathogens (Supplementary note figure 3c). In Supplementary note 3, we analyzed pesticides from all ecosystem groups for further analysis.

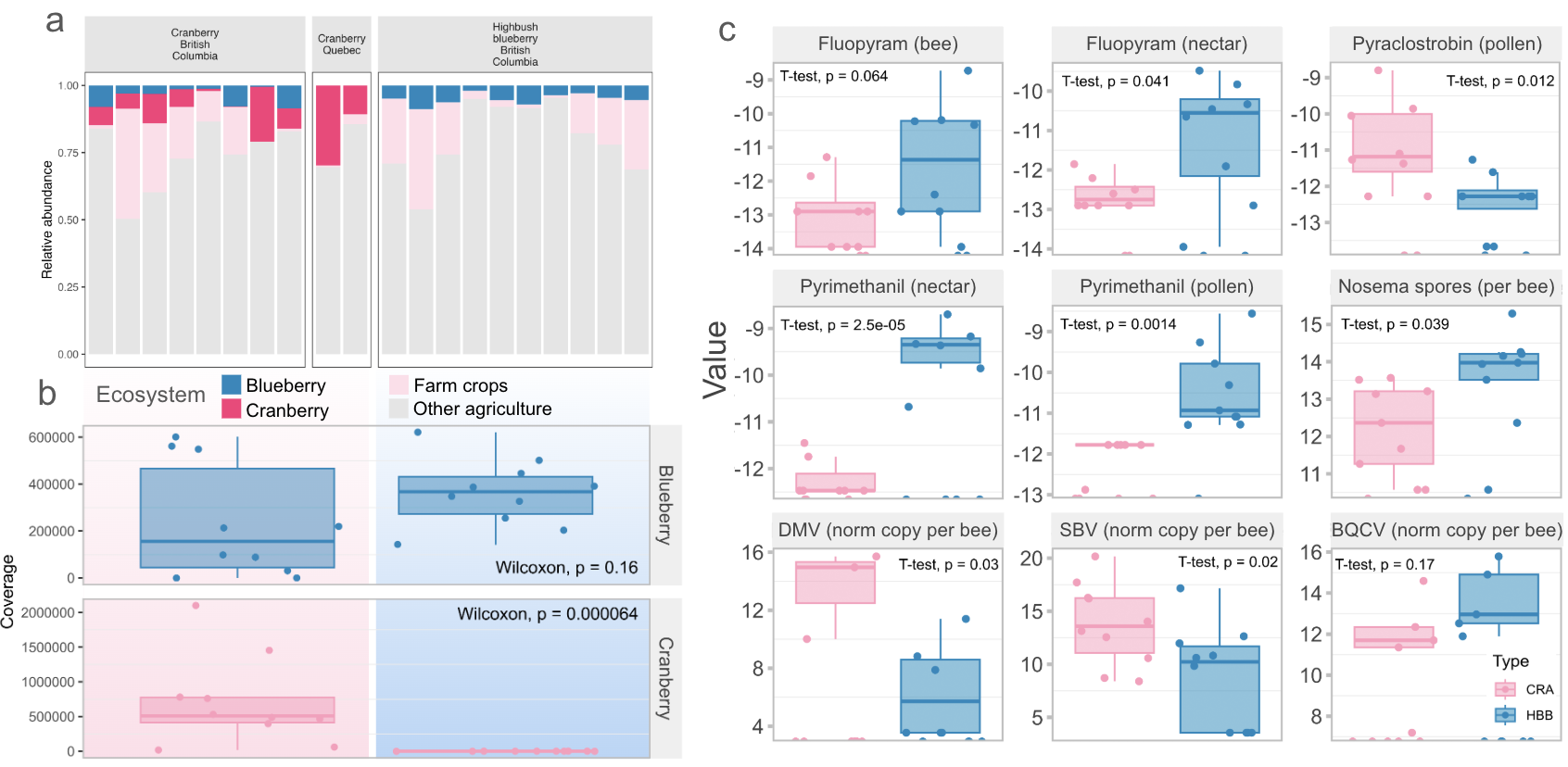

**Supplementary note figure 3.** Floral community composition, environment parameters differences and association bacterial community with it across different agricultural ecosystems. a - Relative abundance of land use types in Vancouver and Quebec Box plot, b - Representation of counts of each crops across each crop ecosystem, c - Representation (log10) of various environmental factors, including pathogens, parasites, pesticide exposure, and other agroecosystem-specific stressors. There were fungicides levels, presented bees, nectar or pollen, Nosema sports on the bees and *A. mellifera* viruses (BQCV, DMV and SBV normalised copies per bee).

Based on the observed differences, we can conclude that the definition of a crop-rich ecosystem is influenced by plant distribution in each area, pesticide use, the presence of viruses and other pathogens and other parameters.

#### Supplementary note 4. Pathogen data analysis

To assess differences in viral infection levels across crop types, we performed within-group analysis of variance using (vegan::betadisper) and between-group analysis of variance using Euclidean distances on log transformed data (vegan::adonis2). Within-group analysis of variance with permutations revealed no significant differences, but between-group analysis of variance revealed that viral variances between different crop ecosystems differed significantly, explaining approximately 60% of the variation in viral load, and PCoA ordination further demonstrated clear separation between crop types (Supplementary note figure 4b). Effect size estimates from linear models (ANOVA) indicated that growing conditions explained more than 35% of the variance for each virus (Supplementary table 5), suggesting a strong influence of crop-related factors on viral infection dynamics in honey bees.

Next we performed collinearity analysis with the stats package. As a result half of the pesticide variables had VIF > 5, indicating notable collinearity with crop (and/or with other pesticides). We also performed fdb-RDA analysis, which shows us that for some crop ecosystems some virus pathogens were more common. It can be seen on the figure 3b and it is seen, that IAPV and VDV viruses were more represented in the CRA and HBB crop ecosystems, and based on the t.test on the log data it was found significant difference in the level of those viruses between HBB, CRA and other crop ecosystems (Supplementary note figure 4a). On the other hand there was a significant difference in the BQCV level in the CRA and HBB to other crops.

We subsequently quantified the extent to which crop ecosystem characteristics and pathogen burden explained variation in the microbiome. As described above, crop environment alone accounted for approximately 36% of the variation in gut microbiome composition. In contrast, a permutation model including only viral pathogen variables explained 21% of the variation (R² = 0.21, p < 0.001). When both crop-related variables and pathogen levels were incorporated into a combined model, the explained variance increased to 44% (R² = 0.44, p < 0.001), with a shared contribution of ~13% between the two predictor groups. This overlap indicates that crop environment and pathogen burden are partially correlated and together shape the microbiome.

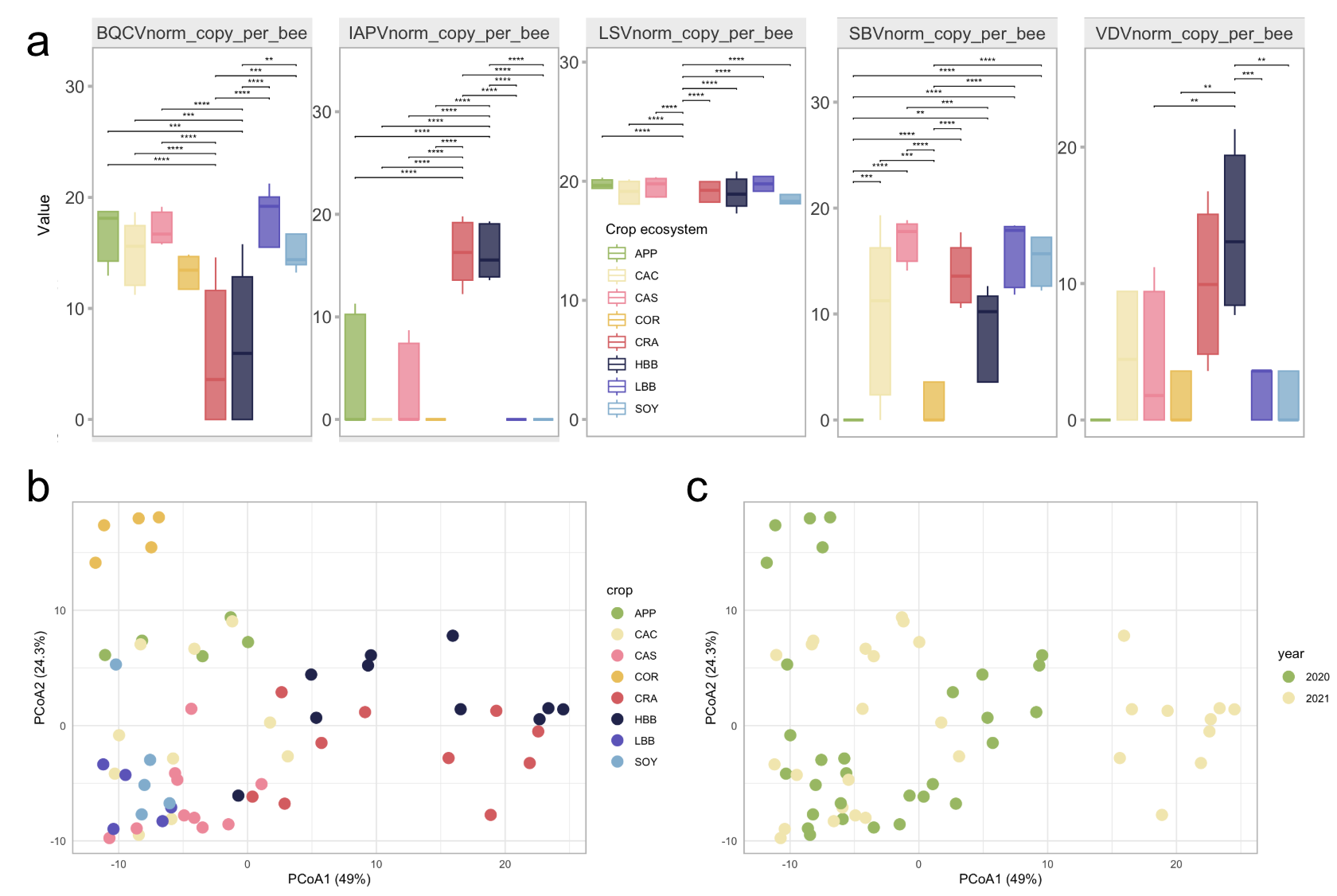

**Supplementary note figure 4. Pathogene variables explanation and connection to the crop ecosystems**

(a) Comparison of log transformed pathogens data (llog(Pathogene+1).). Comparison with pairwise t test.
(b) Principal Coordinate Analysis (PCoA) on Euclidean distances based on pathogens log transformed counts with colors based on crop ecosystem.
(c) Principal Coordinate Analysis (PCoA) on Euclidean distances based on pathogens log transformed counts with colors based on sampled year.

Ultimately, the prevalence of IAPV differed between the CAC and CAS crops in Alberta, despite the close agronomic similarity of these systems. Similarly, SBV abundance varied between SOY and CAC within the same region, and VDV showed differential representation between HBB and CRA crop ecosystems in British Columbia.

To characterize relationships among viral pathogens, we conducted correlation analyses using both Pearson and Spearman coefficients. DWV-B and IAPV displayed a strong positive correlation, and both viruses showed significant negative correlations with BQCV (p < 0.001 for all comparisons; Supplementary note figure 4a). LSV was not detected in any COR crop ecosystem samples, and SBV was absent in the APP ecosystem and occurred at significantly lower levels in COR samples. Neither LSV nor SBV exhibited statistically significant correlations with the other viruses, likely reflecting their restricted presence across ecosystems. Overall, the correlation patterns were consistent with results obtained from the fdb-RDA analysis, reinforcing the conclusion that pathogen interactions and ecosystem context jointly shape viral infection profiles in honey bees.

#### Supplementary note 5. Ancom-BC2 models for pathogene

1. SBV + BQCV + LSV
2. SBV + IAPV + LSV
3. SBV + VDV + LSV

#### Supplementary note 6. Pathogene in connection to the microbiome variation

To verify these associations while accounting for the environmental context, we repeated the ANCOM-BC2 analysis with crop–ecosystem correction after centering pathogen variables across crop environments. This corrected model reproduced the same direction and identity of key associations (Supplementary figures 1-5), confirming that the observed patterns are robust and not solely driven by crop-related variation. Together, these results provide strong evidence for consistent and biologically meaningful links between specific viral infections and bacterial taxa in the honey bee gut microbiome.

We also ran the model for each virus separately for the two ecosystems to examine trends in bacterial abundance within them. We ran the model only on the CRA and HBB ecosystems, as only in these ecosystems were viruses present in more than 50% of samples, allowing us to speculate about their relationship with bacteria. As a result, we observed the same patterns of abundance changes as in the model adjusted for the crop ecosystem (Supplementary figures 1-5). No statistically significant differences were found for any of the viruses, which may be due to the small sample size (10 samples in each).

Overall, we observed exactly the same results in all cases. Only in one case did we see an increase in the abundance of a bacterium in the presence of a virus, although all other analysis methods showed a decrease in the abundance of this bacterium (the abundance of the BQCV virus increased in the CRA crop ecosystem analysis, but not statistically significantly and at a low level). For all other bacteria, the results were consistent.

Furthermore, we observed that for some viruses, bacteria were present that differed in their high LFC levels across all three data types. However, since we did not find a statistically significant result (*p* > 0.05) confirming these differences in any of the analyses, we cannot comment on the accuracy of the data obtained.

#### Supplementary note 7. Pesticide data analysis

To examine the association between pesticides and microbiome, eight pesticides were used: five were detected in pollen (P), two in nectar (N), and one in bee tissue (B) (Supplementary note figure 5b).

To assess whether crop ecosystem structured pesticide profiles, we computed Euclidean distance matrices and determine within-group variances using the betadisper function with permutations which did not show significant differences (*p* value > 0.05) and between-group variances was tested using permutational multivariate (vegan::adonis2) which has shown that crop ecosystem significantly explained 50.3% of the variance in pesticide composition (R² = 0.503). Principal cCordinate Analysis (PCoA) further showed clear separation of crop ecosystem centroids in ordination space (Supplementary note figures 5d,e). In complementary univariate ANOVA/linear models, crop ecosystems also explained >30% of the variance in each pathogen outcome (R² > 0.30; Supplementary table 5).

We next quantified the extent to which crop ecosystem characteristics and pesticide exposure explained variation in the gut microbiome. The crop environment accounted for approximately 36% (R² = 0.36, p < 0.001) of the variation in microbiome composition and a permutation model including only pesticide-related variables explained 29% of the variation (R² = 0.29, p < 0.001). When crop-related variables and pesticide measures were incorporated into a combined model, the proportion of explained variance increased to 49% (R² = 0.49, p < 0.001), with an estimated ~16% shared contribution between the two predictor sets. This shared variance indicates partial collinearity between crop environment and pesticide burden, suggesting that these factors jointly influence microbiome structure. Taken together, these results support the conclusion that variation in honey bee gut microbiota across crop ecosystems arises from the combined effects of crop environment, pesticide pressure, and agricultural chemical exposure.

As a result, concentrations of Fluopyram in both pollen (p) and nectar (n) were significantly higher in the CAS crop ecosystem. Moreover, several pesticides, including Spirotetramat (p), Metconazole (b), Fluopyram (n and p), and Fenhexamid (p), were predominantly detected in CAS, HBB, and CRA. In contrast, certain compounds were more characteristic of other crop systems; for instance, pyraclostrobin was most abundant in SOY, whereas Boscalid (b) was detected across all crop ecosystems but with substantial variability in concentration.

To characterize relationships among pesticides, we produced correlation analyses using both Pearson and Spearman coefficients. As a result, a strong correlation was found for a number of pesticides. Interestingly, the same pesticide present in pollen and nectar was not always well correlated. For example, Fluopyram (N) and (P) were significantly less correlated with each other than Fluopyram (N) was with Fenhexamide (P).

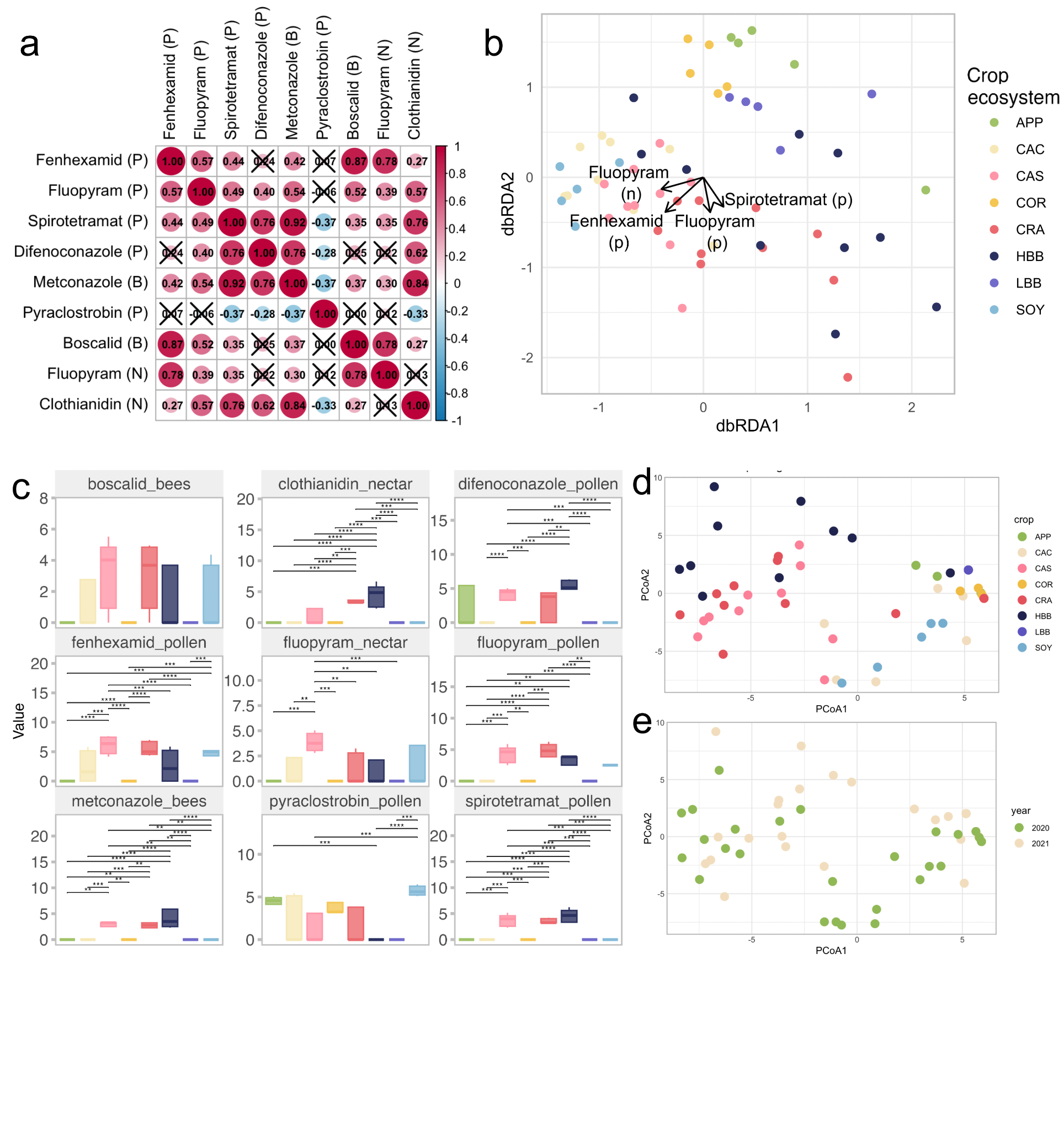

**Supplementary note figure 5. Pesticides variables explanation and connection to the crop ecosystems**

(a) Correlation plot for pathogenic variables with each other based on the Spearman index. X means not significant correlation.
(b) Fdb-RDA ordination plots display microbial community composition constrained by pathogenic variables. Vectors are targeted at crops in which viruses are present in high numbers.
(c) Comparison of log transformed pesticide data (log((Pesticide * 10^7)+1)). Comparison with pairwise t test.
(d) Principal Coordinate Analysis (PCoA) on Euclidean distances based on pesticides log transformed counts with colors based on crop ecosystem.

Principal Coordinate Analysis (PCoA) on Euclidean distances based on pesticides log transformed counts with colors based on sampled year.

#### Supplementary note 8. Ancom-BC2 models for pesticide

1. fenhexamid(p) + difenoconazole(p) + pyraclostrobin(p) + clothianidin(n)
2. fluopyram(p) + pyraclostrobin(p0
3. spirotetramat(p) + pyraclostrobin(p) + boscalid(b) + fluopyram(n)
4. fenhexamid(p) + difenoconazole(p) + pyraclostrobin(p) + boscalid(b) + fluopyram(n)
5. metconazole(b) + pyraclostrobin(p) + boscalid(b) + fluopyram(n)
6. fenhexamid(p) + fluopyram(p) + spirotetramat(p) + difenoconazole(p) + metconazole(b) + pyraclostrobin(p) + boscalid(b) + fluopyram(n) +clothianidin(n)
7. spirotetramat(p) + difenoconazole(p) + metconazole(b) + pyraclostrobin(p) + boscalid(b) +clothianidin(n)
8. spirotetramat(p) + difenoconazole(p) + metconazole(b) + pyraclostrobin(p) + fluopyram(n) +clothianidin(n)
9. fenhexamid(p) + pyraclostrobin(p) + boscalid(b) + fluopyram(n) +clothianidin(n)

#### Supplementary note 9. Pesticides in connection to the microbiome variation

To confirm trends in bacterial abundance in samples related to pesticide use, an ANCOM analysis was conducted. To monitor the persistence of population trends, three data sets were analyzed to assess the repeatability of the experiment. The experiment was repeated using data adjusted for the influence of the culture ecosystem after centering pathogen variables based on crop growth conditions and on two separate ecosystems (CRA and HBB) (Supplementary figures 8-16).

We generally observed repeatability of the trends obtained in the main results. Moreover, we often observed that bacteria statistically significantly associated with pesticide parameters in the main results also had a relatively strong effect in the replicated results. For example, for boscalid (b), clothianidin (n), fluopyram (n), and fluopyram (p), the effect was quite strong and consistent with the original results (Supplementary figures 8,9,12,13 respectively). Fluopyram (n) and *Apibacter* sp. wkB309 demonstrated the strongest consistent effect, with LFC exceeding 0.5 in all cases. For clothianidin (n), fluopyram (n), and fluopyram (p) and their associated bacteria, the effect sizes typically ranged from 0.2 to 0.6, which, despite the scatter, was consistent with the original results (with a LFC of 0.4 and statistical significance).

However, for some pesticides, the opposite effect was demonstrated, and the association between the pesticides and the bacterial organisms was not confirmed. For example, for metconazole (b) and spirotetromat (p), the effect sizes were negative in some assays (Supplementary figures 14, 16). This may indicate inconsistency in the data and a false association previously identified.

We also found other statistically significantly associated bacteria for some pesticides in our analysis. For example, a statistically significant association was found for fenhexamid (p) in the centered, adjusted data and in the CRA analysis (with *Apibacter* sp. wkB309 and *Bartonella apis*, respectively) (Supplementary figures 11). But this observed association was not replicated in the CRA analysis of the data. In fact, an inverse association was found for this pesticide with these bacterial organisms.

A similar pattern was observed for difenoconazole (p), which had a strong negative association with *Spiroplasma melliferum* (statistically significant in the CBD data) (Supplementary figures 10). However, difenoconazole (p) had the opposite (positive) effect when analyzed separately in the CRA data.

We also observed a strong association between pesticides and pathogens in some cases, although statistical significance was not determined. For example, we observed a strong negative effect (LFC -0.5 or -1) for boscalid (b) against *Mellisococcus plutonius* (Supplementary figures 8). Although the effect was not statistically significant, we can still hypothesize that there is some correlation between the high abundance of this bacterium and the use of this pesticide.

### SUPPLEMENTARY FIGURES

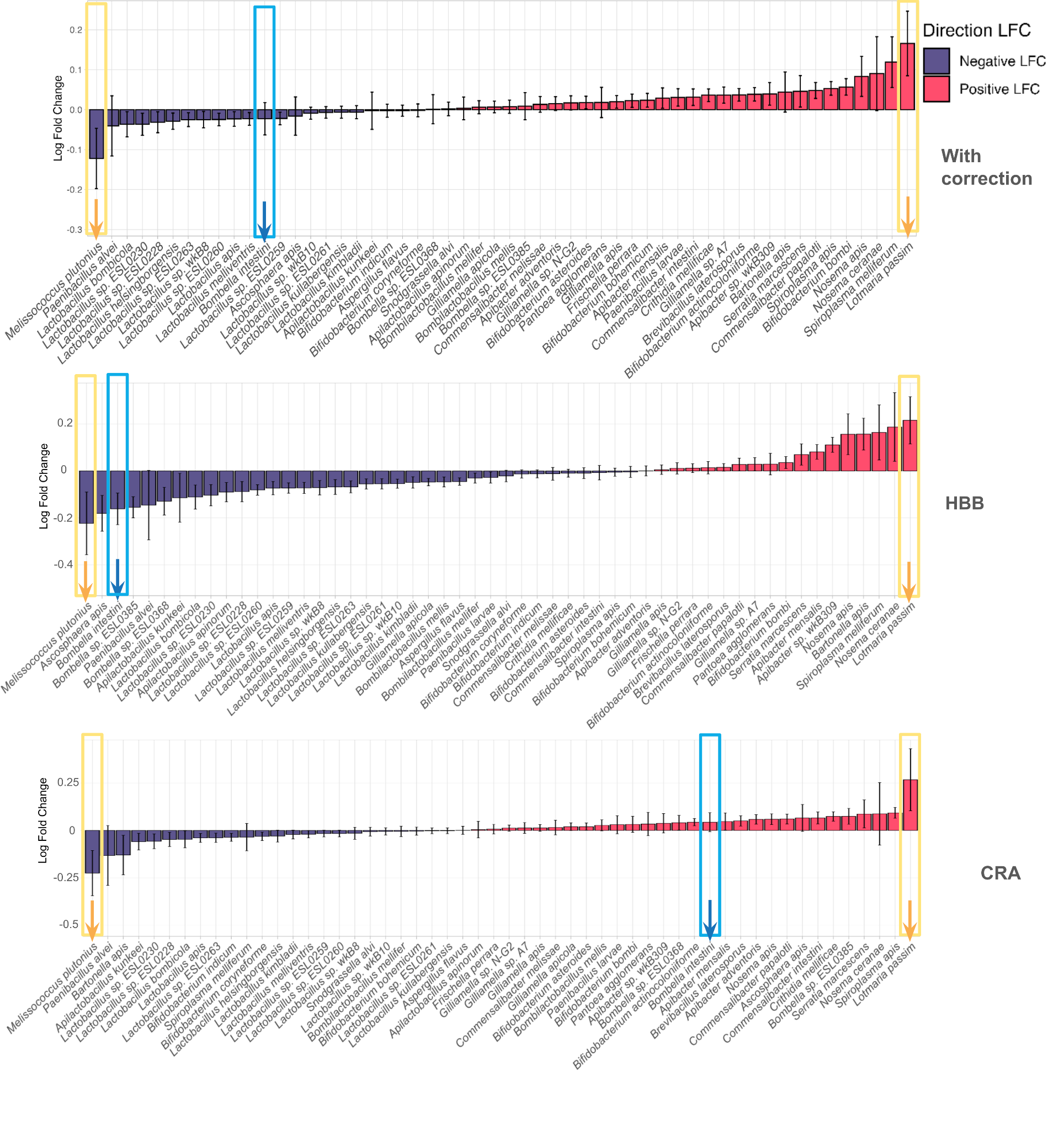

**Supplementary figure 1.** Log fold change (LFC) of species abundance between presented and not presented BQCV virus from (a) all samples and years with centration of viruses and crop inclusion, (b) only from HBB sample and (c) only from CRA samples. Statistical significance (ANCOM-BC2; p <0.05) is indicated by *. Positive LFC indicates an increase in abundance between factors, presence compared to absence for each virus pathogen, whereas negative LFC indicates a decrease.

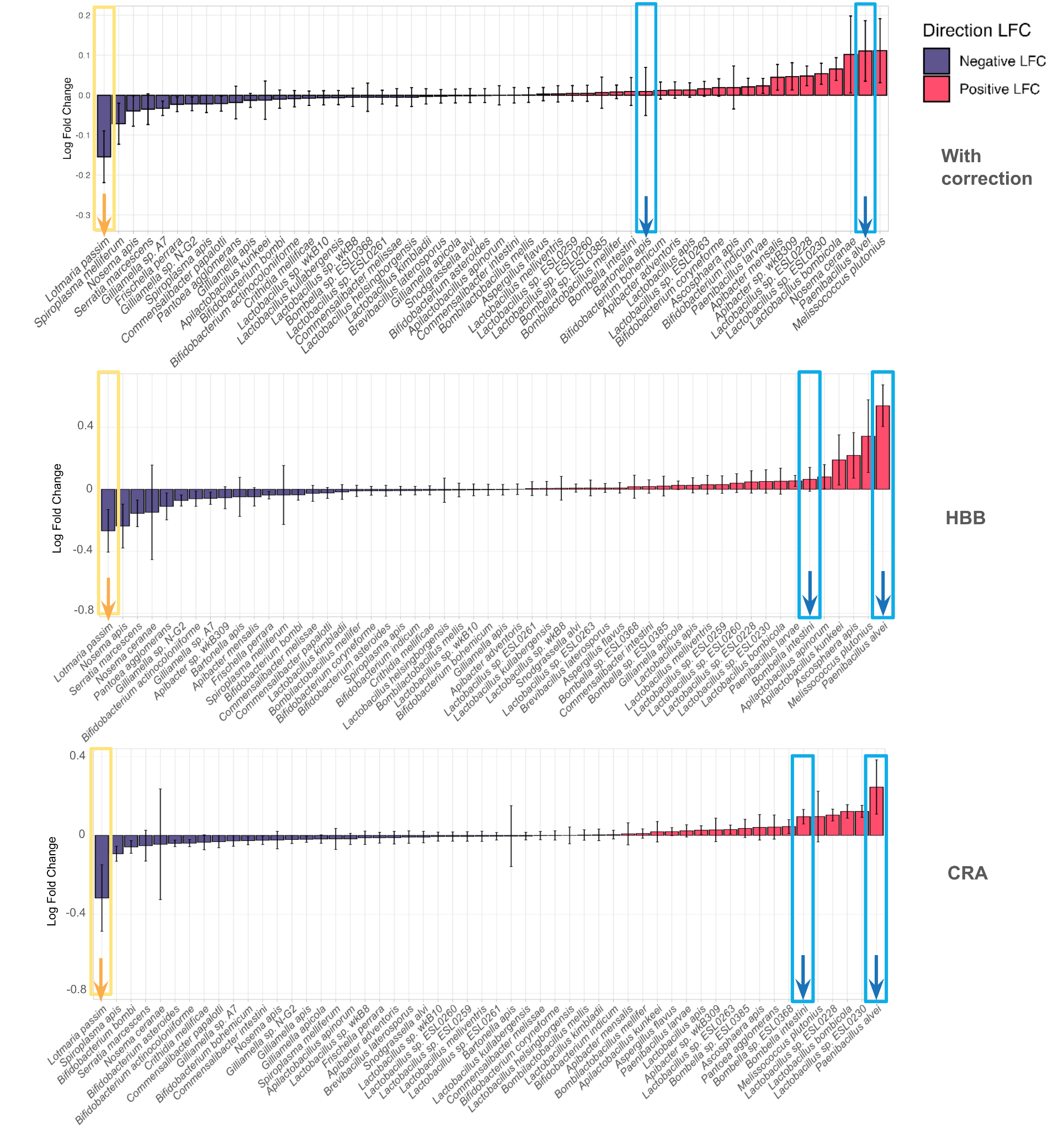

**Supplementary figure 2.** Log fold change (LFC) of species abundance between presented and not presented IAPV virus from (a) all samples and years with centration of viruses and crop inclusion, (b) only from HBB sample and (c) only from CRA samples. Statistical significance (ANCOM-BC2; p <0.05) is indicated by *. Positive LFC indicates an increase in abundance between factors, presence compared to absence for each virus pathogen, whereas negative LFC indicates a decrease.

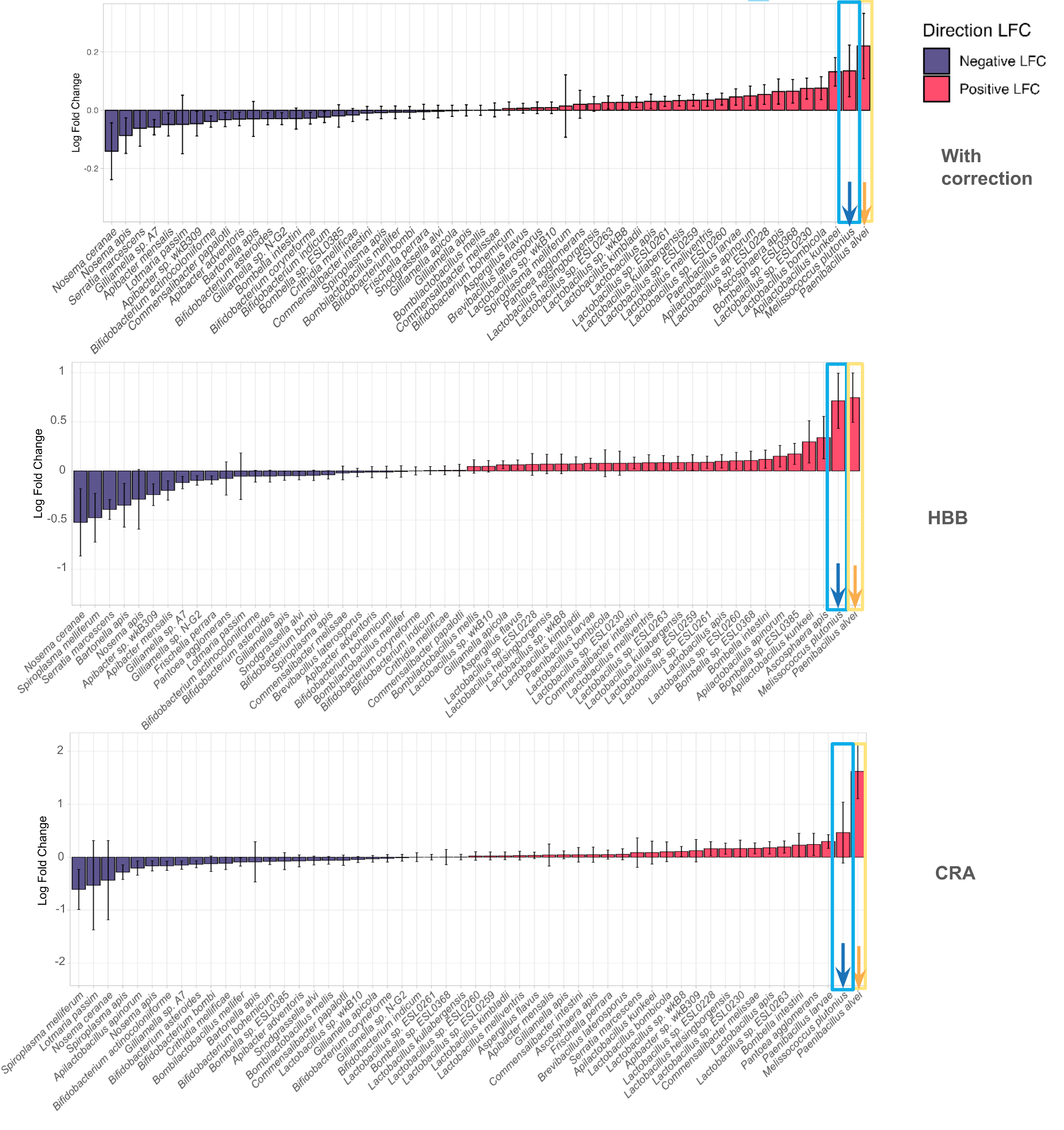

**Supplementary figure 3.** Log fold change (LFC) of species abundance between presented and not presented LSV virus from (a) all samples and years with centration of viruses and crop inclusion, (b) only from HBB sample and (c) only from CRA samples. Statistical significance (ANCOM-BC2; p <0.05) is indicated by *. Positive LFC indicates an increase in abundance between factors, presence compared to absence for each virus pathogen, whereas negative LFC indicates a decrease.

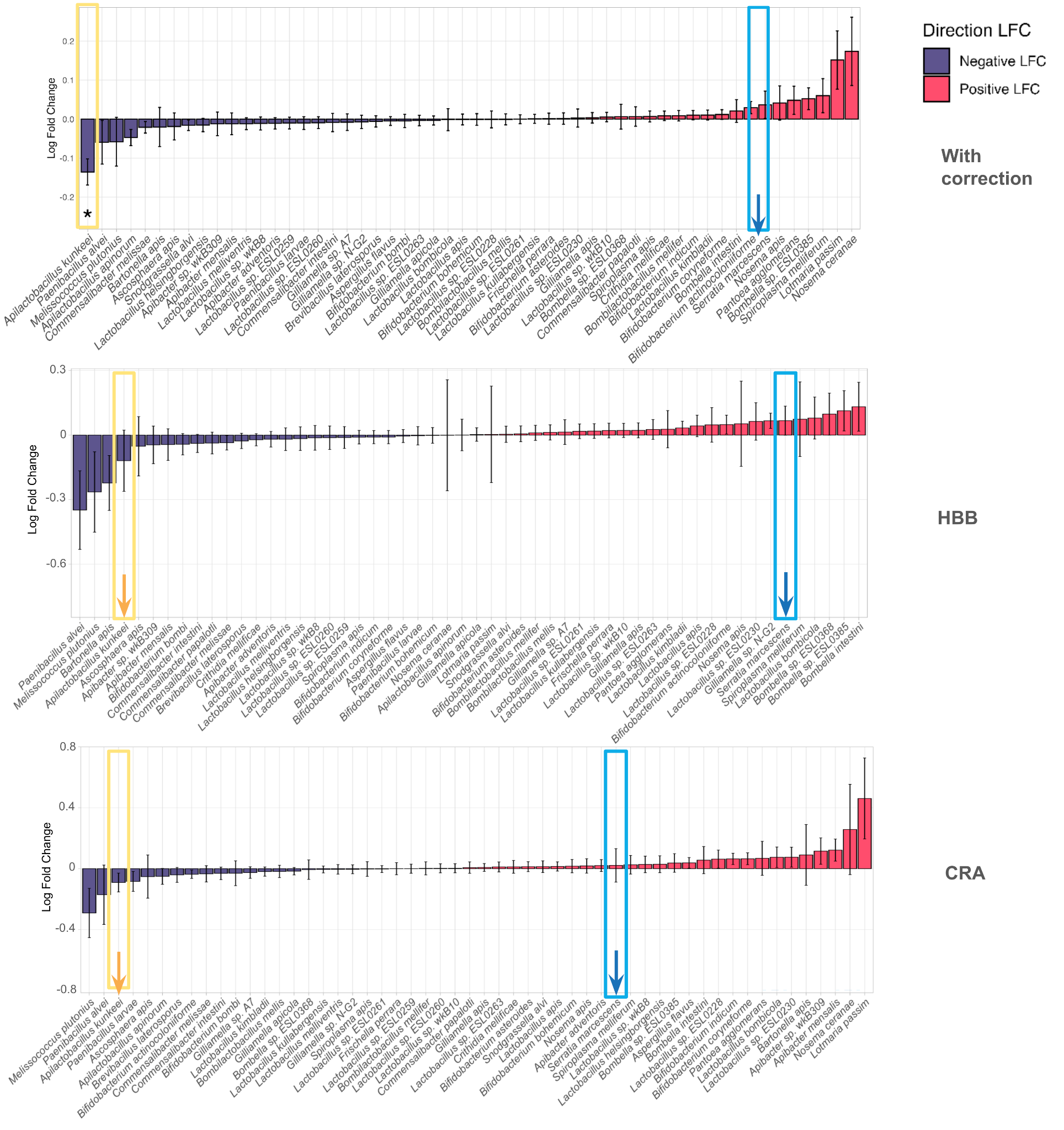

**Supplementary figure 4.** Log fold change (LFC) of species abundance between presented and not presented SBV virus from (a) all samples and years with centration of viruses and crop inclusion, (b) only from HBB sample and (c) only from CRA samples. Statistical significance (ANCOM-BC2; p <0.05) is indicated by *. Positive LFC indicates an increase in abundance between factors, presence compared to absence for each virus pathogen, whereas negative LFC indicates a decrease.

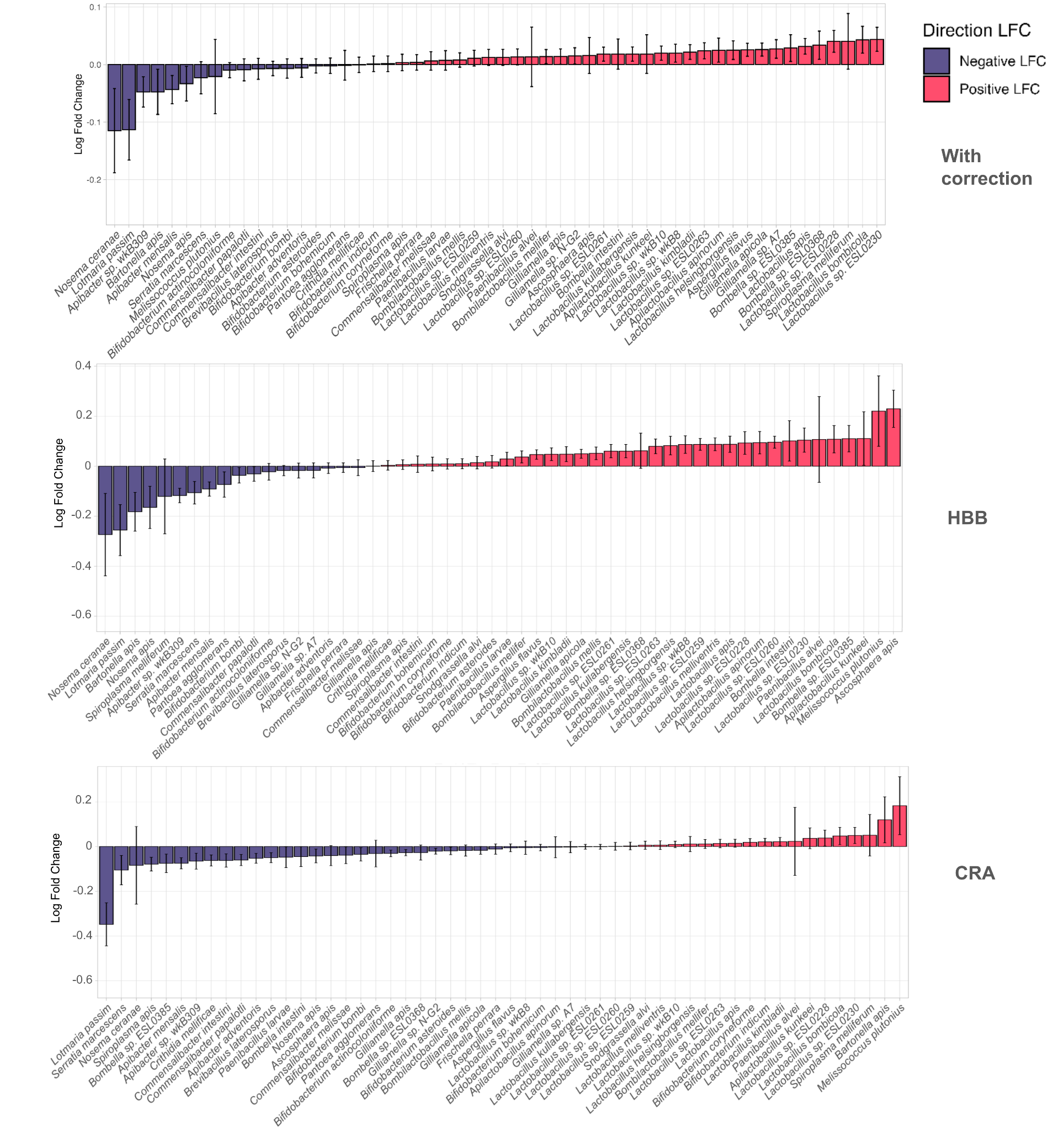

**Supplementary figure 5.** Log fold change (LFC) of species abundance between presented and not presented VDV virus from (a) all samples and years with centration of viruses and crop inclusion, (b) only from HBB sample and (c) only from CRA samples. Statistical significance (ANCOM-BC2; p <0.05) is indicated by *. Positive LFC indicates an increase in abundance between factors, presence compared to absence for each virus pathogen, whereas negative LFC indicates a decrease.

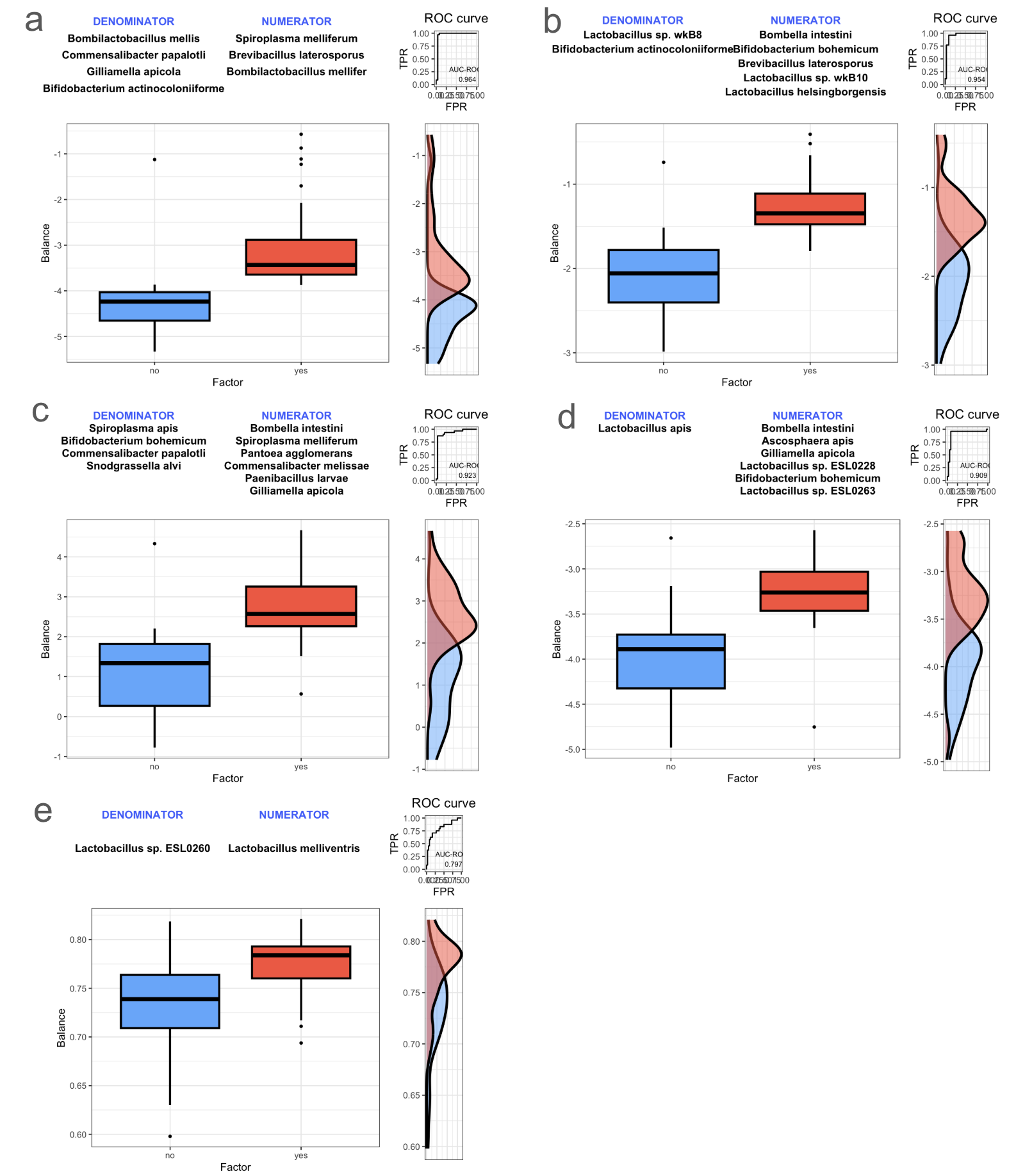

**Supplementary figure 6.** SelBal analysis with ROC AUC for pesticides from pollen: (а) fenhexamid, (b) spirotetromat, (c) fluopyram, (d) difenoconazole, (e) pyraclostrobin.

Each boxplot shows the SelBal balance comparing samples in which the pesticide is absent (“No”) versus present (“Yes”). If the “Yes” group displays higher balance values than the “No” group, this indicates that the numerator taxa are relatively more abundant (or the denominator taxa less abundant) in pesticide-positive samples.

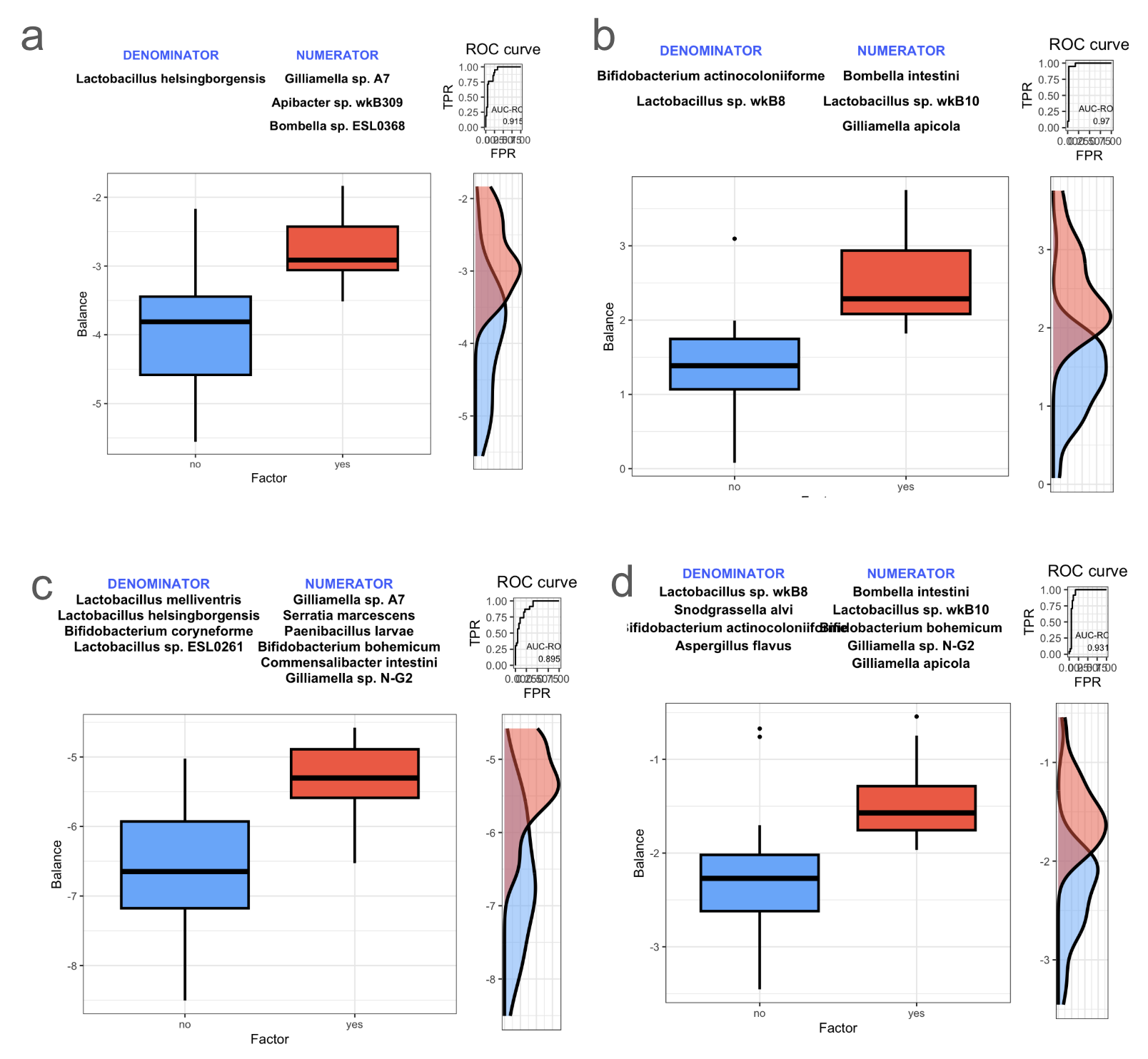

**Supplementary figure 7.** SelBal analysis with ROC AUC for pesticides from nectar and bees: (а) fluopyram (n), (b) clothianidin (n), (c) boscalid (b), (d) metconazol (b).

Each boxplot shows the SelBal balance comparing samples in which the pesticide is absent (“No”) versus present (“Yes”). If the “Yes” group displays higher balance values than the “No” group, this indicates that the numerator taxa are relatively more abundant (or the denominator taxa less abundant) in pesticide-positive samples.

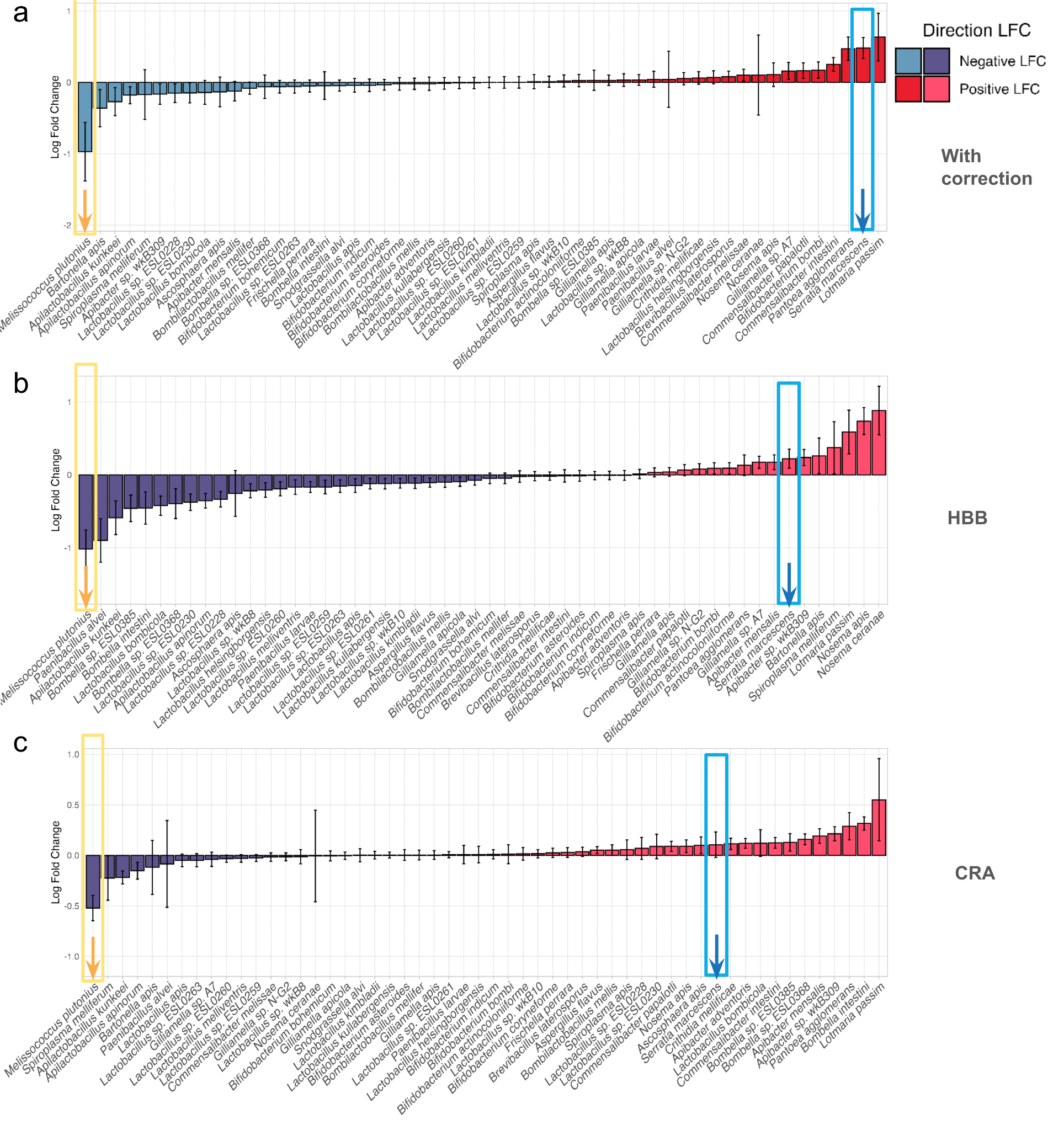

**Supplementary figure 8.** Log fold change (LFC) of species abundance between presented and not presented boscalid (b) pesticide from (a) all samples and years with centration of viruses and crop inclusion, (b) only from HBB sample and (c) only from CRA samples. Statistical significance (ANCOM-BC2; p <0.05) is indicated by *. Positive LFC indicates an increase in abundance between factors, presence compared to absence for each virus pathogen, whereas negative LFC indicates a decrease.

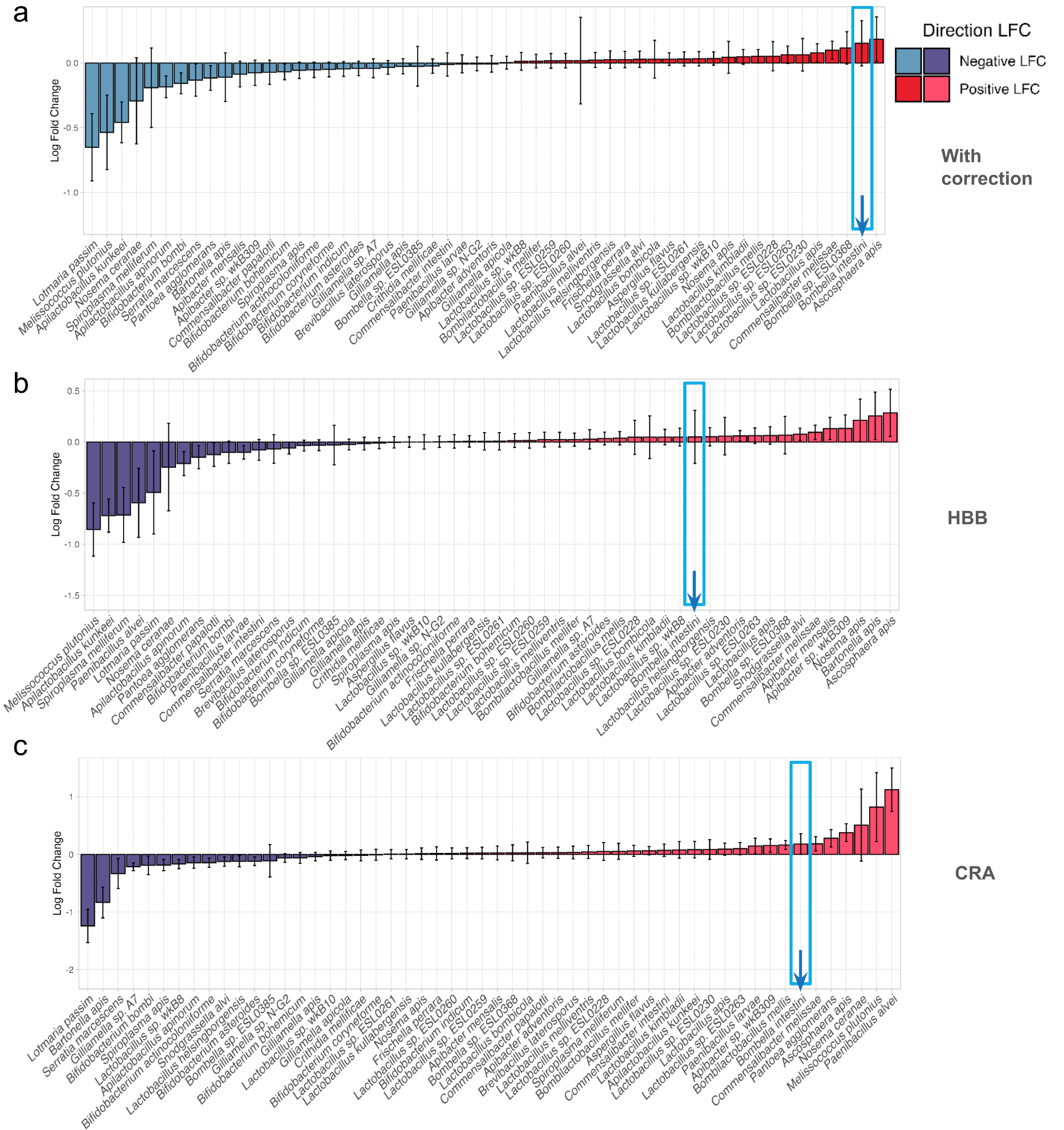

**Supplementary figure 9.** Log fold change (LFC) of species abundance between presented and not presented clothianidin (n) pesticide from (a) all samples and years with centration of viruses and crop inclusion, (b) only from HBB sample and (c) only from CRA samples. Statistical significance (ANCOM-BC2; p <0.05) is indicated by *. Positive LFC indicates an increase in abundance between factors, presence compared to absence for each virus pathogen, whereas negative LFC indicates a decrease.

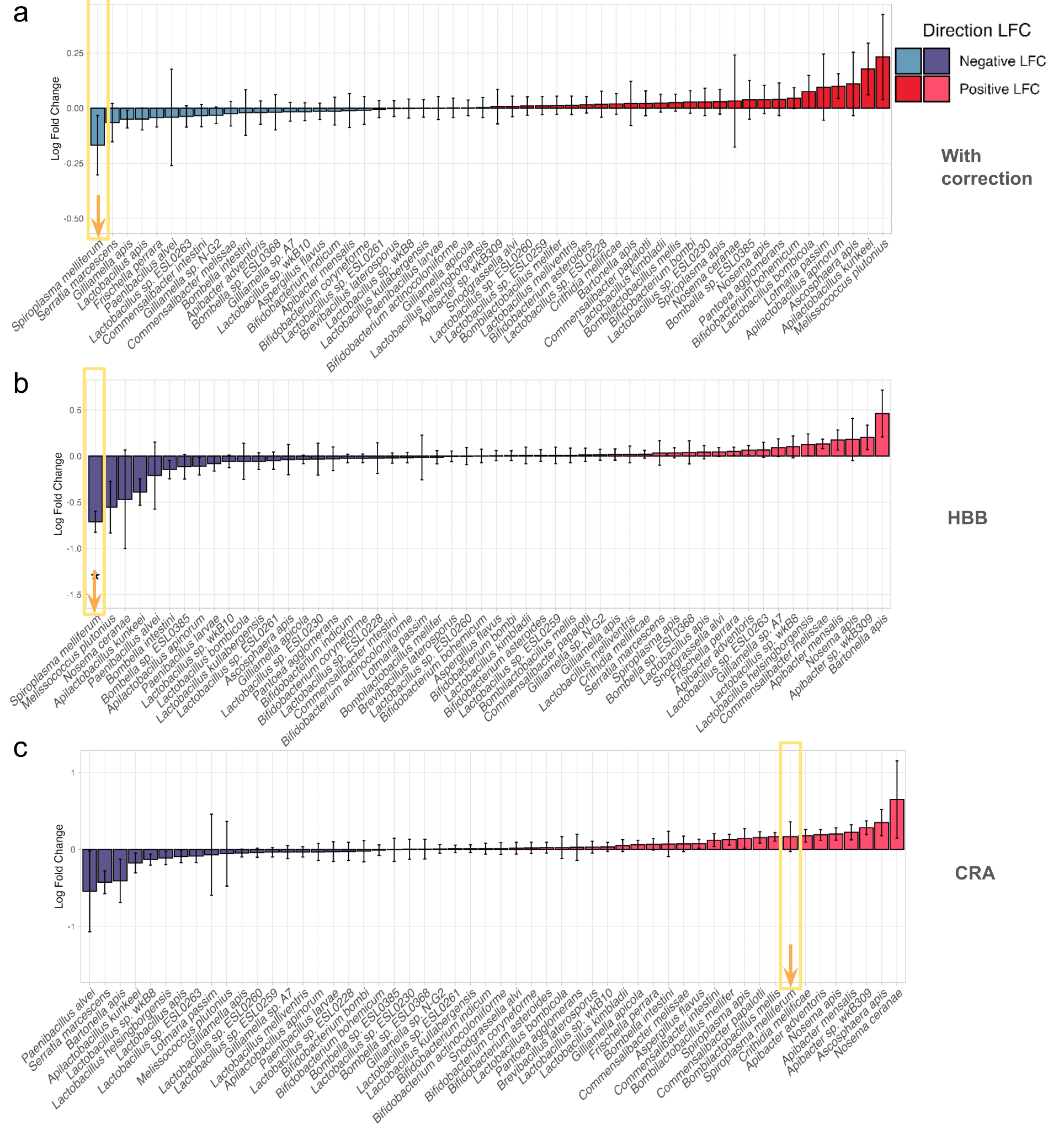

**Supplementary figure 10.** Log fold change (LFC) of species abundance between presented and not presented difenoconazole (p) pesticide from (a) all samples and years with centration of viruses and crop inclusion, (b) only from HBB sample and (c) only from CRA samples. Statistical significance (ANCOM-BC2; p <0.05) is indicated by *. Positive LFC indicates an increase in abundance between factors, presence compared to absence for each virus pathogen, whereas negative LFC indicates a decrease.

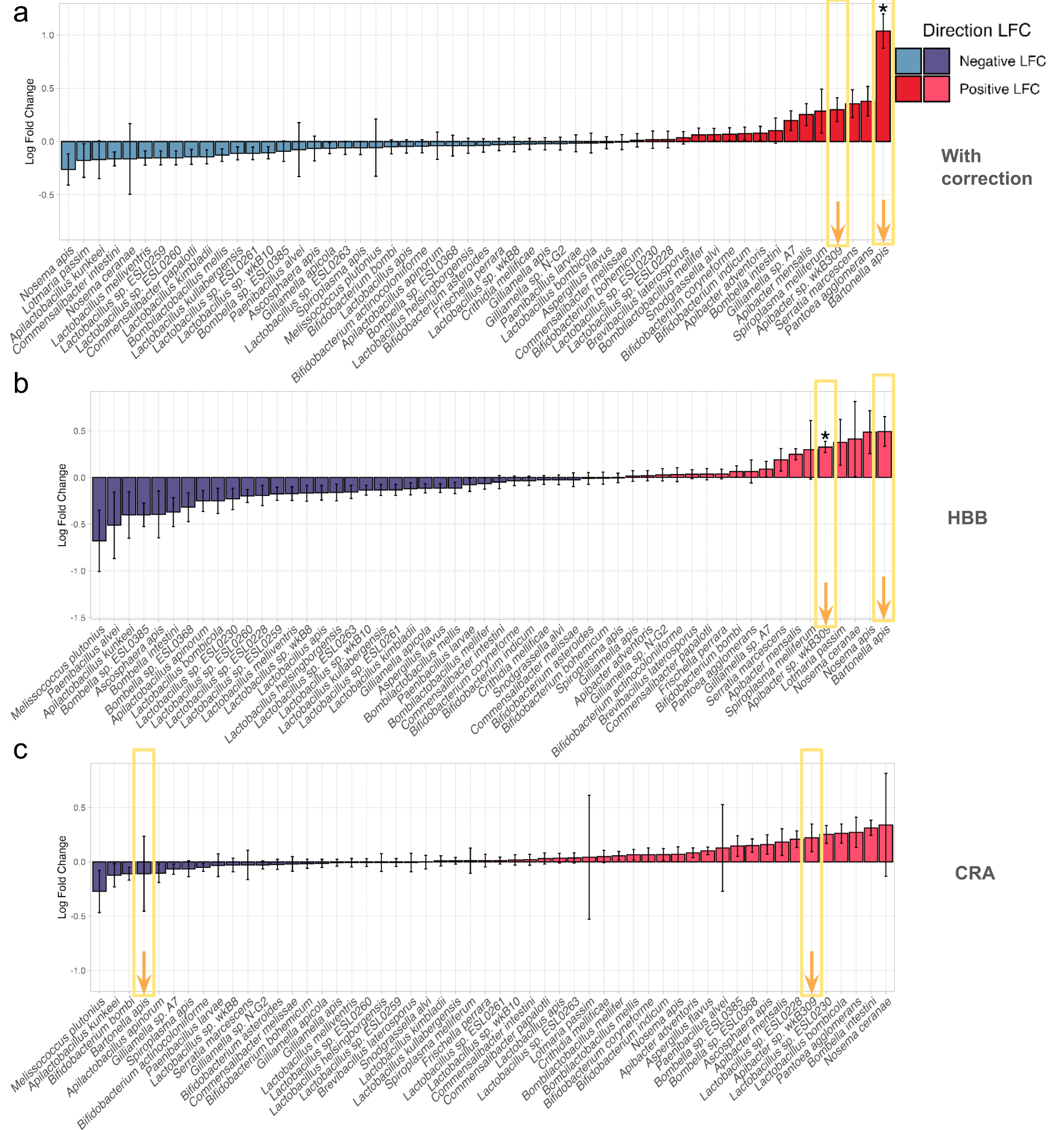

**Supplementary figure 11.** Log fold change (LFC) of species abundance between presented and not presented fenhexamid (p) pesticide from (a) all samples and years with centration of viruses and crop inclusion, (b) only from HBB sample and (c) only from CRA samples. Statistical significance (ANCOM-BC2; p <0.05) is indicated by *. Positive LFC indicates an increase in abundance between factors, presence compared to absence for each virus pathogen, whereas negative LFC indicates a decrease.

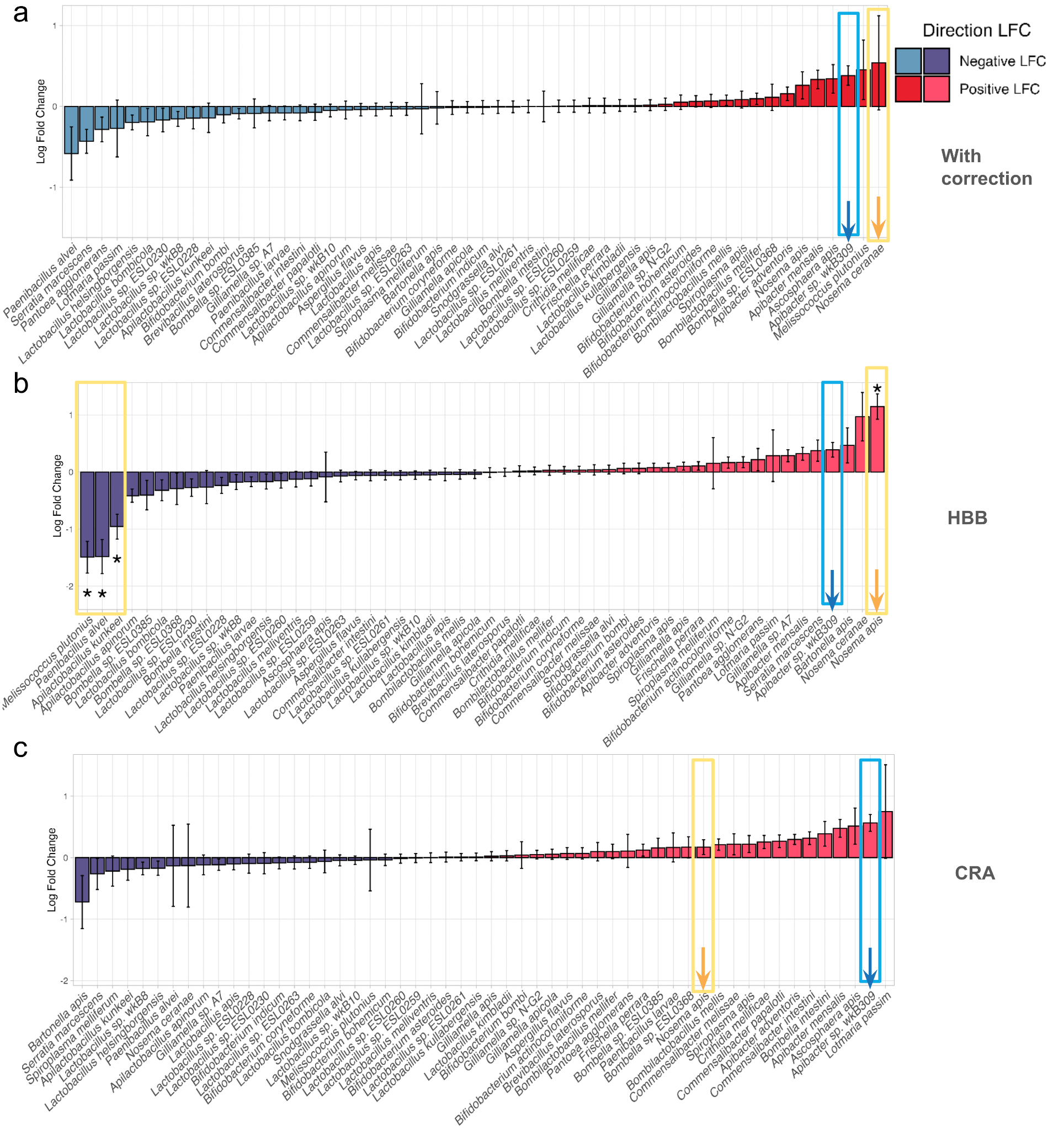

**Supplementary figure 12.** Log fold change (LFC) of species abundance between presented and not presented fluopyram (n) pesticide from (a) all samples and years with centration of viruses and crop inclusion, (b) only from HBB sample and (c) only from CRA samples. Statistical significance (ANCOM-BC2; p <0.05) is indicated by *. Positive LFC indicates an increase in abundance between factors, presence compared to absence for each virus pathogen, whereas negative LFC indicates a decrease.

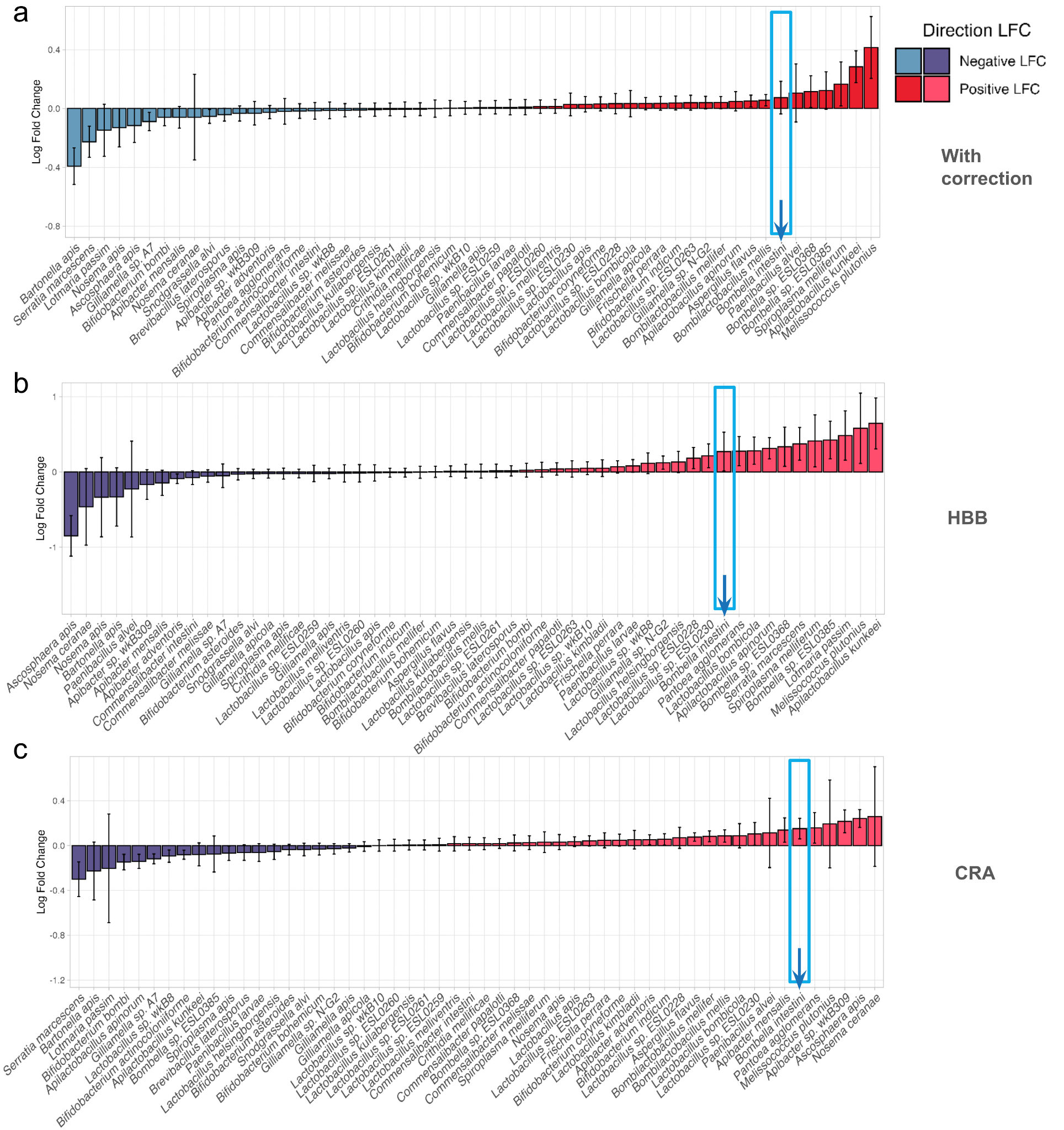

**Supplementary figure 13.** Log fold change (LFC) of species abundance between presented and not presented fluopyram (p) pesticide from (a) all samples and years with centration of viruses and crop inclusion, (b) only from HBB sample and (c) only from CRA samples. Statistical significance (ANCOM-BC2; p <0.05) is indicated by *. Positive LFC indicates an increase in abundance between factors, presence compared to absence for each virus pathogen, whereas negative LFC indicates a decrease.

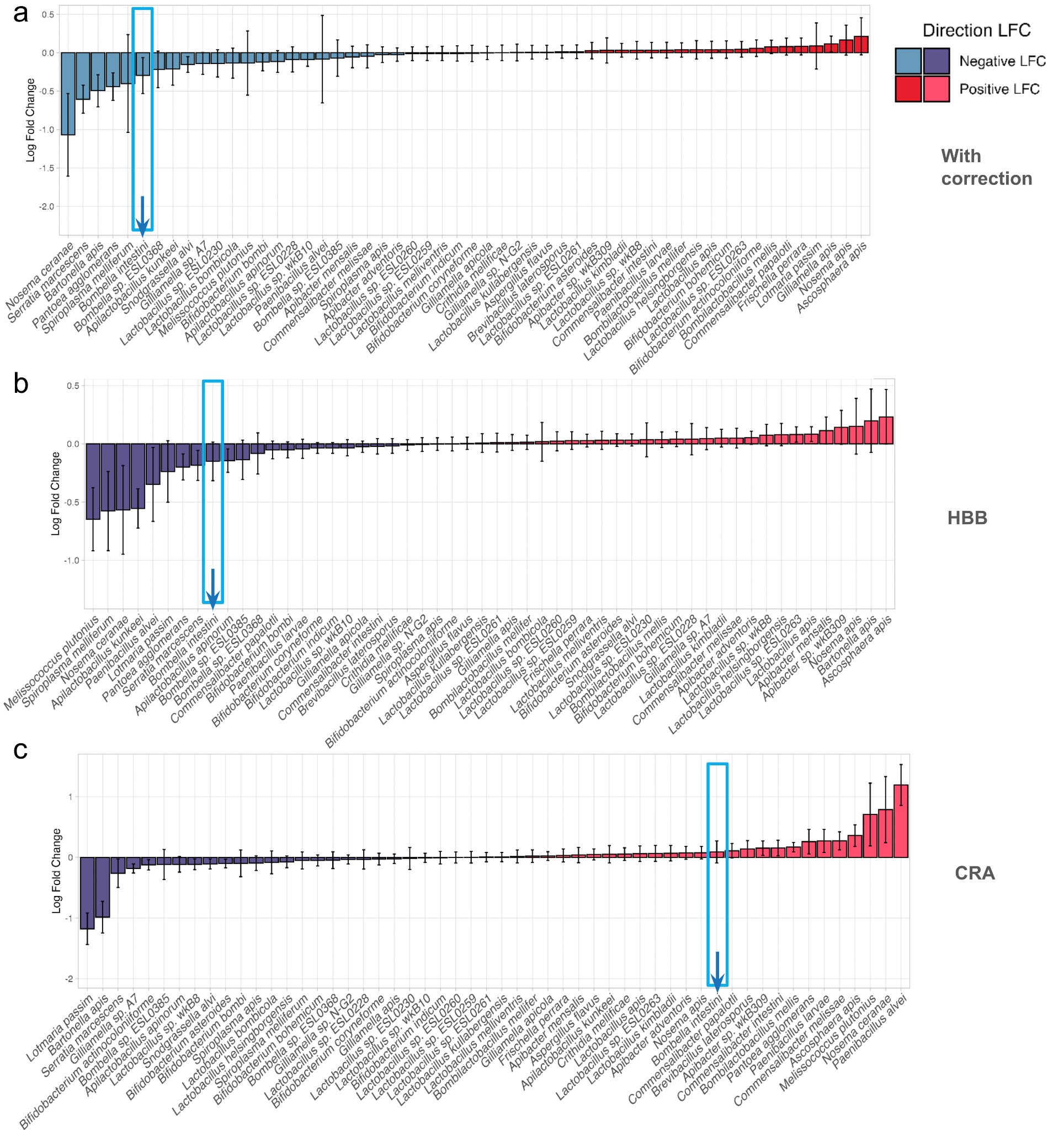

**Supplementary figure 14.** Log fold change (LFC) of species abundance between presented and not presented metconazole (b) pesticide from (a) all samples and years with centration of viruses and crop inclusion, (b) only from HBB sample and (c) only from CRA samples. Statistical significance (ANCOM-BC2; p <0.05) is indicated by *. Positive LFC indicates an increase in abundance between factors, presence compared to absence for each virus pathogen, whereas negative LFC indicates a decrease.

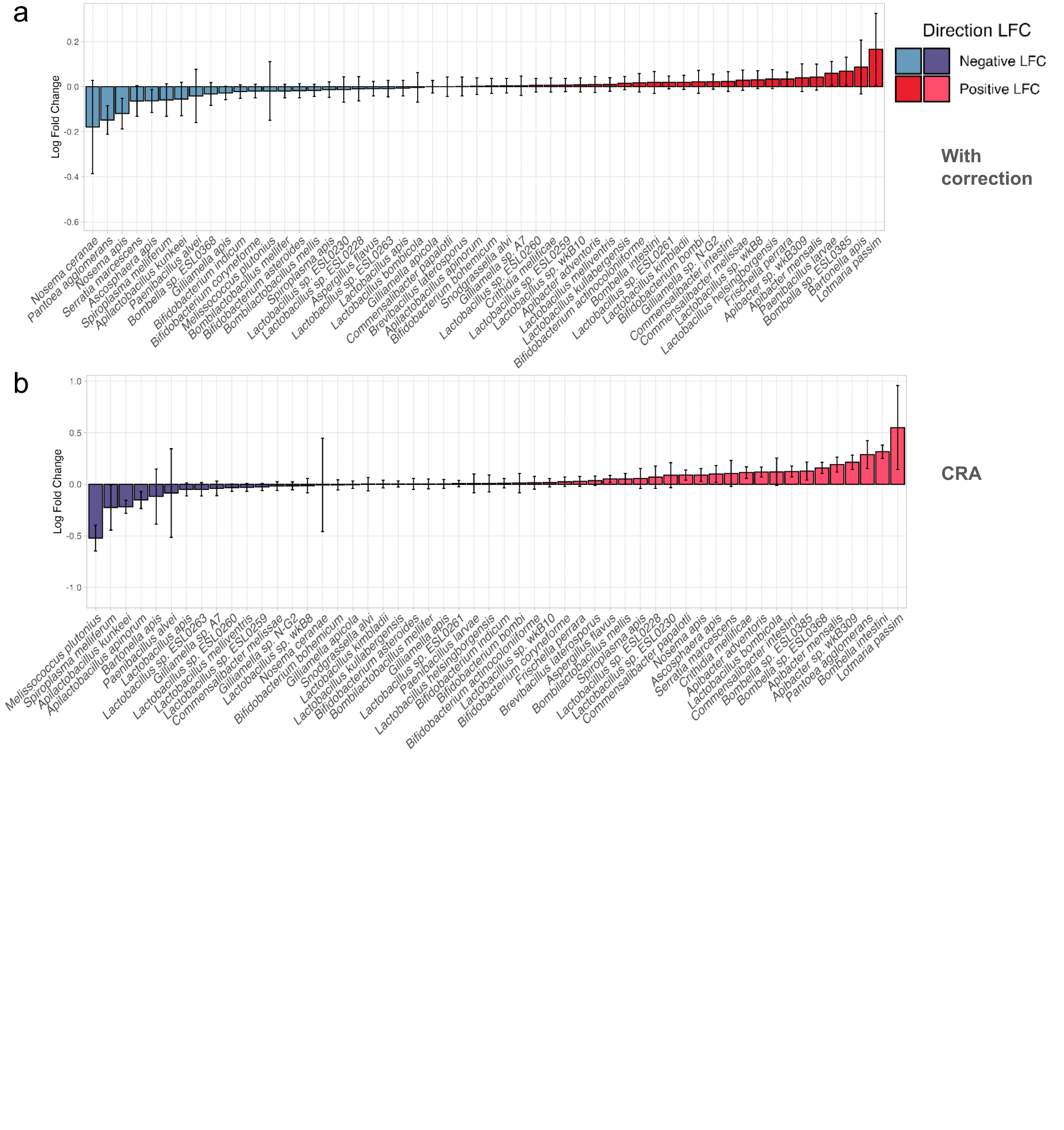

**Supplementary figure 15.** Log fold change (LFC) of species abundance between presented and not presented pyraclostrobin (p) pesticide from (a) all samples and years with centration of viruses and crop inclusion, (b) only from HBB sample and (c) only from CRA samples. Statistical significance (ANCOM-BC2; p <0.05) is indicated by *. Positive LFC indicates an increase in abundance between factors, presence compared to absence for each virus pathogen, whereas negative LFC indicates a decrease.

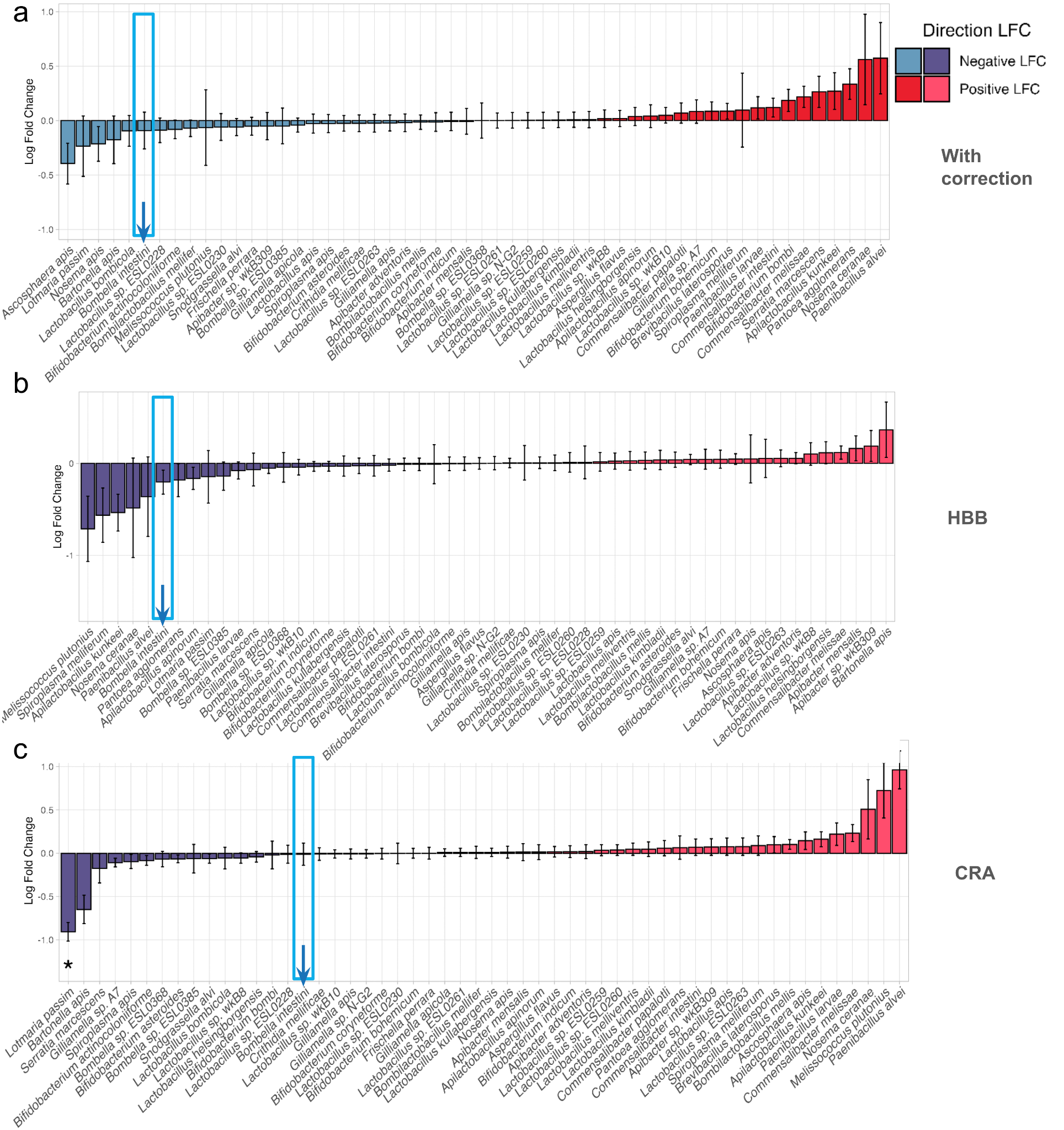

**Supplementary figure 16.** Log fold change (LFC) of species abundance between presented and not presented spirotetromat (p) pesticide from (a) all samples and years with centration of viruses and crop inclusion, (b) only from HBB sample and (c) only from CRA samples. Statistical significance (ANCOM-BC2; p <0.05) is indicated by *. Positive LFC indicates an increase in abundance between factors, presence compared to absence for each virus pathogen, whereas negative LFC indicates a decrease.
